# X-Y divergence of house fly (*Musca domestica*) proto-sex chromosomes follow distinct evolutionary trajectories despite residing in the same genome

**DOI:** 10.64898/2026.09.26.754726

**Authors:** Jae Hak Son, David Luecke, Basanta Bista, Yesbol Manat, Weihuan Cao, Leo Beukeboom, Daniel Bopp, Christopher E Ellison, Perot Saelao, Richard P Meisel

## Abstract

Sex determination systems and sex chromosomes frequently differ between species. One cause of these differences is new sex determining genes that drive evolutionary turnover of sex chromosomes. At the earliest stages of this turnover, X and Y (or Z and W) chromosomes start out as nearly identical homologs, and they can diverge via chromosomal rearrangements (e.g., inversions) that suppress X-Y recombination. However, multiple examples from across animals and plants provide exceptions to this canonical model of sex chromosome evolution. For example, some X-Y pairs remain undifferentiated for long evolutionary time periods, while other sex chromosomes become differentiated without chromosomal rearrangements. How or why these non-canonical trajectories occur remains elusive, despite increasing evidence of their pervasiveness. The house fly, *Musca domestica*, is a well-suited system to address this gap because all six chromosomes can be a Y, providing multiple replicates of a natural experiment within a single genomic environment. To test for canonical and non-canonical evolutionary trajectories, we generated haplotype-resolved chromosome-level assemblies from five strains of the house fly, each of which carries a different Y chromosome (I^M^, II^M^, III^M^, V^M^, and Y^M^). We identified an inversion on only one of the sex chromosomes (II^M^), which was associated with elevated X-Y divergence but did not capture the male-determining locus. In contrast, there was X-Y divergence across almost the entire length of the I^M^, III^M^, and V^M^ sex chromosomes, despite no detectable inversions. Y^M^ was the only sex chromosome to contain substantial Y-specific sequences, which were limited to a segment on one end of the chromosome containing the male-determining gene. This Y^M^ chromosome, and its corresponding X, were highly diverged from the X chromosome found in many other flies (Muller element F), despite a strong cytological resemblance. This study highlights how multiple different canonical and non-canonical modes of sex chromosome evolution can co-exist within a single genome.

## Introduction

Sex chromosomes and sex determination can evolve rapidly, even though sex determination is an essential developmental process [1,2]. The rapid evolution of sex chromosomes can include fusions with autosomes, forming neo-sex-chromosomes. In addition, an autosome can acquire a new sex-determining locus, becoming a proto-sex-chromosome. After a neo- or proto-sex chromosome arises, the new X and Y (or Z and W) chromosomes can become differentiated from each other [3]. This X-Y differentiation is thought to be promoted by chromosomal rearrangements, such as inversions, that suppress X-Y recombination. Suppressed X-Y recombination is hypothesized to maintain genetic linkage between the sex-determining locus and sexually antagonistic (male-beneficial and female-deleterious) alleles on the Y chromosome [4,5]. This model therefore predicts that loci with the tightest genetic linkage to the sex-determiner should have the highest levels of X-Y differentiation at the early stages of sex chromosome evolution. Subsequent inversions and other chromosomal rearrangements can drive additional X-Y differentiation, creating strata of differing divergence times [6]. A key feature of this X-Y divergence is the degeneration of the Y chromosome, including pseudogenization of genes and the accumulation of repetitive elements [7].

Despite the theoretical expectations regarding X-Y divergence, the evolution of sex chromosomes often does not follow the predicted trajectory. For example, if there is no male recombination genome-wide, the inherent lack of genetic exchange in males can allow for X-Y differentiation in the absence of sex-chromosome-specific suppressors of recombination. Such is the case in *Drosophila*, where male achiasmy leads to the pseudogenization of many genes along the entire length of young neo-Y chromosomes within a very short evolutionary time [8,9]. In contrast, European tree frogs have male heterochiasmy (i.e., crossing over), but males undergo genetic recombination within very limited regions of chromosomes [10]. The majority of the Y chromosome, which does not recombine with the X, can therefore diverge from the X, stratifying X-Y divergence along the lengths of the sex chromosomes without the requirement of inversions [11]. However, X-Y differentiation in some frogs can be reverted by transient X-Y recombination in sex-reversed XY females, maintaining undifferentiated (homomorphic) sex chromosomes for long evolutionary time periods [12–14]. Similarly, paleognathous birds (e.g., kiwis, cassowaries, emus, rheas, and ostriches) have retained homomorphic ZW chromosomes for > 100 million years (My) through Z-W recombination over large chromosomal regions, which differs from the highly degenerated W chromosomes of neognathous birds [15,16]. Reconciling these non-canonical evolutionary trajectories with the conventional model of sex chromosome evolution is an important yet unresolved goal in evolutionary genomics [14,17,18].

Brachyceran flies (*i.e.*, higher Diptera) are a well-suited model system in which to contrast canonical and non-canonical sex chromosome evolution because there are variable rates of sex chromosome turnover and differentiation among lineages [19]. The ancestral karyotype in Brachycera is considered to be a set of six chromosome pairs (2n=12), consisting of five gene-rich euchromatic autosomal pairs (Muller elements A-E) and a gene-poor heterochromatic X chromosome (Muller element F) [20–23]. The ancestral male karyotype has a gene-poor Y chromosome with a male-determining locus, while females have two X chromosomes [19,24].

This karyotype and sex chromosome system appears to be retained in many families, including Tephritidae (true fruit flies) and Calliphoridae (blow flies) [24,25]. Across Brachycera, synteny (i.e., gene content) is well conserved within the Muller elements [19–21]. However, chromosomal inversions are common within elements [26,27]. In addition, element A became a proto-X chromosome before the common ancestor of the *Drosophila* genus, and element F transitioned to an autosome [22]. Within *Drosophila*, there have been multiple fusions of element A and its corresponding Y chromosome with other elements, creating independent neo-sex chromosomes in various *Drosophila* species [28]. Furthermore, in the family Muscidae, the ancestral karyotype of six pairs of chromosomes appears to be retained in many lineages [29], but the heterochromatic X chromosome does not appear to be homologous to element F [30,31]. In Muscidae species, element F has been lost from the karyotype, [30–32], and one of the other elements appears to have become a proto- or neo-sex chromosome [32].

Among the Muscidae, the house fly, *Musca domestica*, is especially well-suited to study sex chromosome evolution because it has multiple very young proto-sex chromosomes. The *Musca domestica male determiner* (*Mdmd*) gene has been found on four different chromosomes (II, III, V, and Y), each of which was transformed into a proto-Y with the addition of *Mdmd* [33]. In addition, chromosome I can also carry a male-determining locus, transforming it into a proto-Y, but the male-determiner is not *Mdmd* and is not yet identified [33,34]. The homologous, non-male-determining chromosome in a given male genotype is, by default, a proto-X for that particular genotype. We refer to the male-determining (*M*) loci on each chromosome as *M*^I^, *M*^II^, *M*^III^, *M*^V^, and *M*^Y^; the proto-Y chromosomes are referred to as I^M^, II^M^, III^M^, V^M^, and Y^M^; and flies carrying a specific proto-Y chromosome are referred to as M1, M2, M3, M5, and MY males [35]. The *M*^II^ and *M*^III^ loci are near the centromeres on euchromatic chromosomes [35], suggesting that the house fly male-determiners arose in pre-existing low recombining regions. In contrast to the other proto-Y chromosomes in house fly, Y^M^ is a gene-poor, heterochromatic chromosome, much like a canonical Y. Unlike a canonical Y chromosome, Y^M^ likely arose from the gene-poor and heterochromatic X chromosome, meaning that it began its evolutionary trajectory gene-poor and heterochromatic, instead of evolving to that state in the canonical path [36]. Moreover, chromosome IV can be a Z/W sex chromosome if it carries a female-determining allele of *transformer* (*Md-tra^D^*), which is epistatically dominant and desensitized to *Mdmd* activity [37].

Females with the *Md-tra^D^* allele can therefore carry one or more proto-Y chromosomes [38]. The house fly polygenic sex determination system thus provides multiple proto-sex chromosomes that resemble and differ from the canonical trajectories in unique ways within a single genomic environment. However, detailed examination of these evolutionary trajectories has been historically limited due to the lack of technologies that could generate phased chromosome-level assemblies.

In this study, we generated phased chromosome-scale assemblies from males of five house fly strains that each carries a different proto-Y chromosome (I^M^, II^M^, III^M^, V^M^, or Y^M^). This allowed us to characterize the very early, and possibly non-canonical, evolution of Y chromosomes. These different young sex chromosomes evolved in the same genomic environment, effectively creating a repeated natural experiment to test hypotheses about Y chromosome evolution. We evaluated the differences in DNA sequence, chromosome structure, and repeat content between the proto-Y and proto-X chromosomes in order to identify canonical and non-canonical evolutionary trajectories.

## Results

### Chromosome-scale Genome Assemblies

We generated highly contiguous male genome assemblies from five different strains of house flies using PacBio HiFi reads and Hi-C reads. Each strain harbors a *M* locus on a different chromosome, with five of the six chromosomes represented (Table 1). For the best contiguity, we first constructed primary assemblies of the five strains, which switch between two haplotypes to bridge the sequencing gaps. The final assembly of most of the strains contained six large scaffolds corresponding to the six chromosomes (Table 1). The Hi-C contact map showed a high intensity of intrachromosomal interactions within scaffolds, indicating that each chromosome was assembled into each scaffold in each strain (Figure 1, Figure S1). Although the scaffolds are technically pseudo-chromosomes [39], we hereafter refer to the six scaffolds as house fly chromosomes. We also constructed two haplotype-resolved (phased) assemblies for each strain, which we used to estimate heterozygosity within strains and to examine polymorphic inversions (see below). For strains that are heterozygous for a polymorphic inversion, we resolved primary assemblies by prioritizing haplotypes that are structurally co- linear with the chromosomes in the other strains (Figure S1).

**Figure 1.**
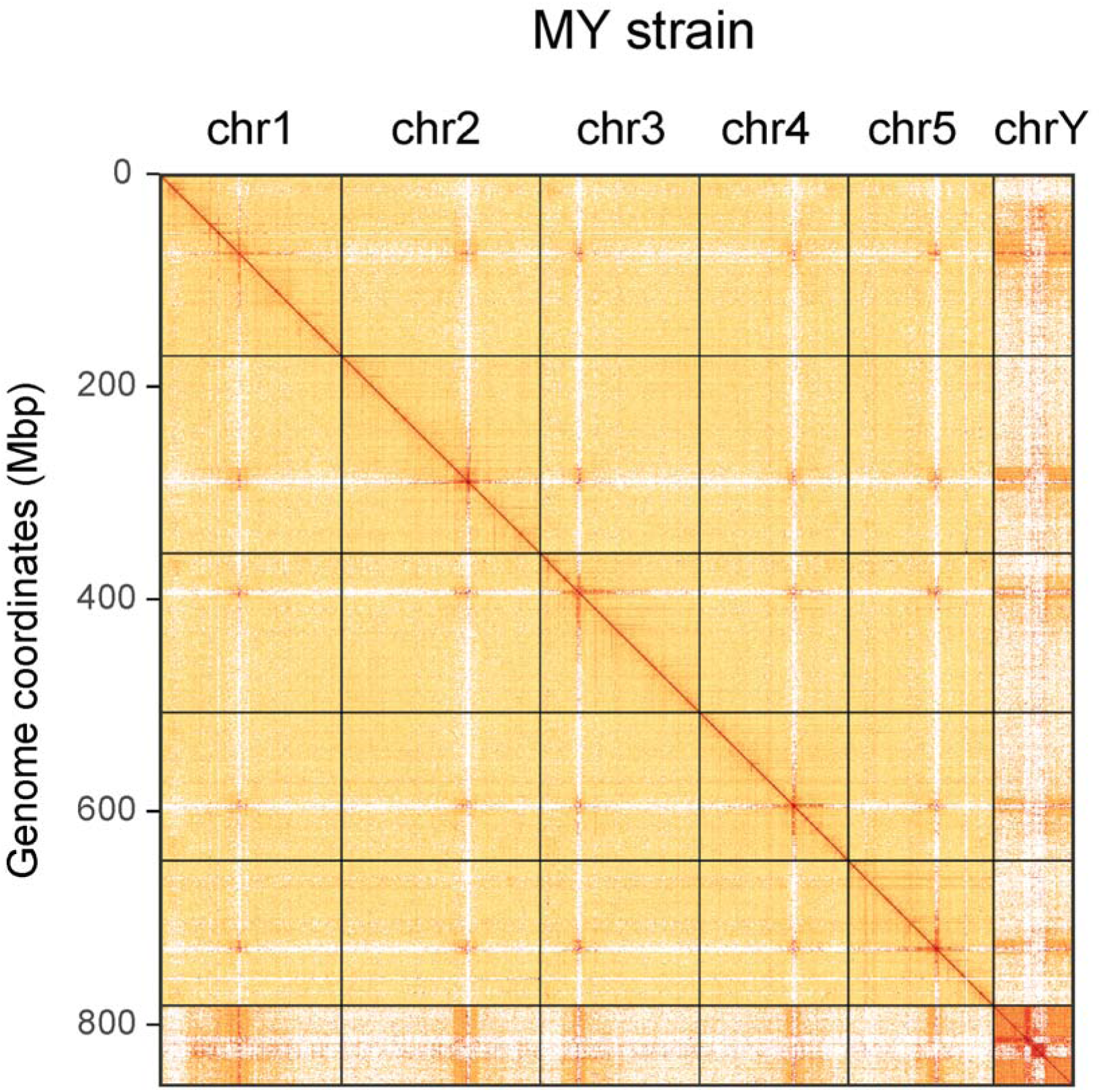
HiC contact map from six chromosomes (I-V, and Y) of the MY strain.

**Table 1.**
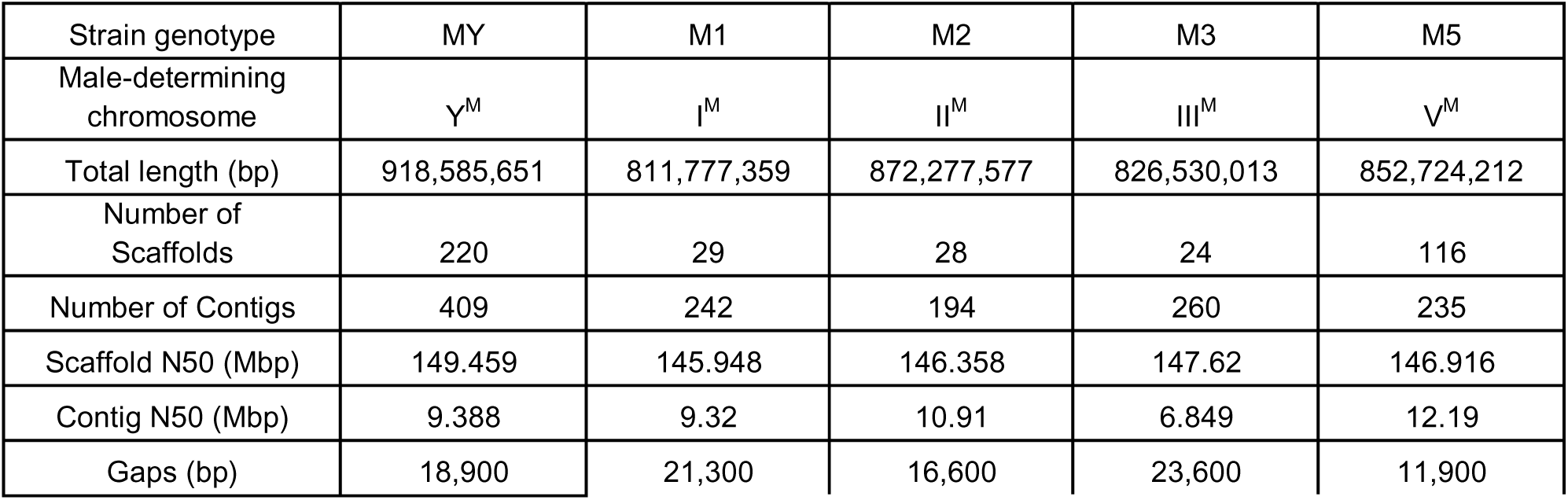
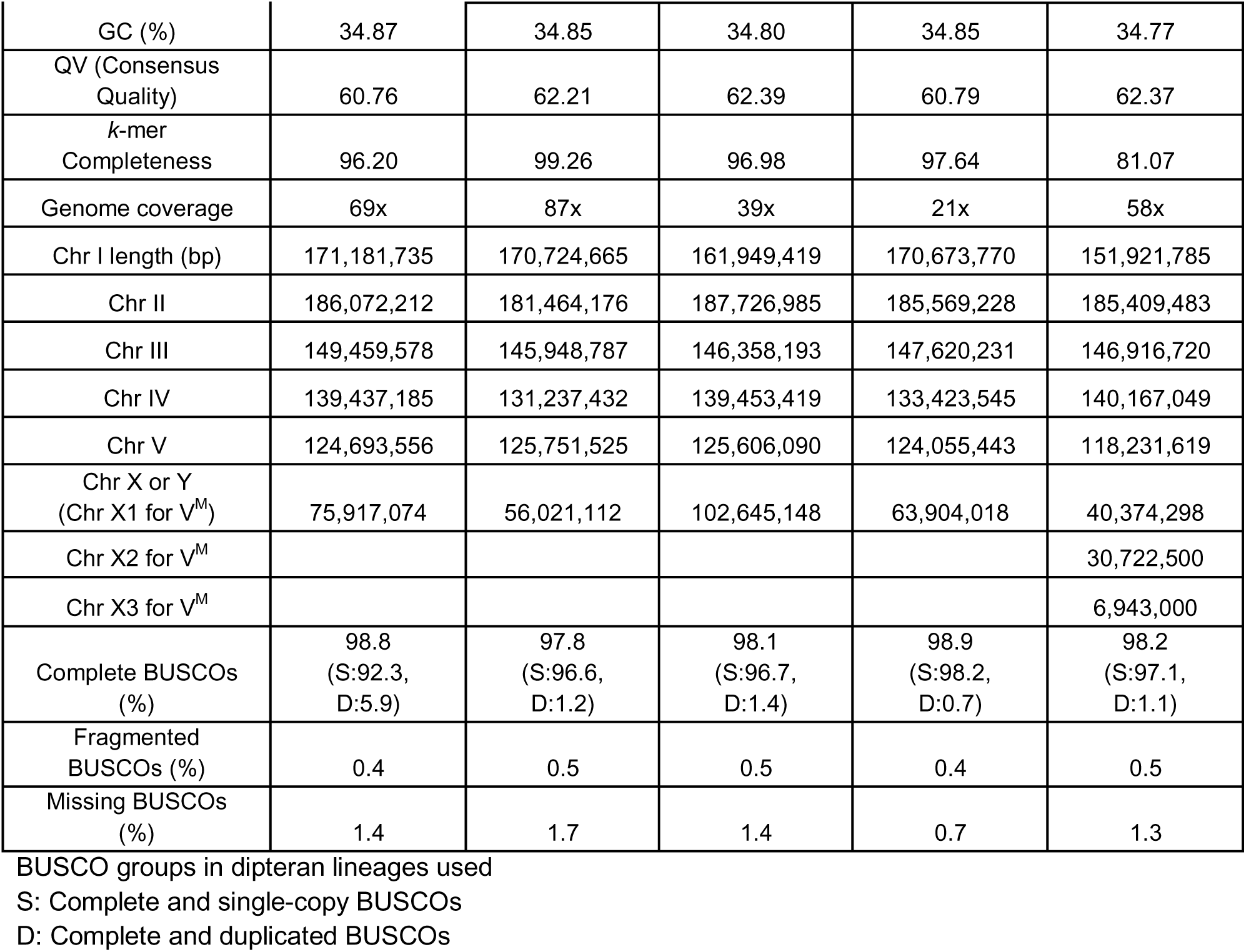
Summary of statistic and BUCSO analyses for the five strain genome assemblies

The sixth scaffold proved to be the most challenging to assemble across all genomes. This sixth scaffold likely corresponds to the X chromosome in most strains (see below). However, in the M5 strain, it appears that the sixth, seventh, and eighth largest scaffolds belong to separate regions of the X chromosome (Table 1). We could not determine the order and orientation of the scaffolds to generate a single scaffold representing the entire X chromosome in the M5 strain because the intrachromosomal interactions among the three scaffolds did not clearly appear in the Hi-C contact map (Figure S1). In addition, in the MY strain primary assembly, the sixth scaffold contains the male-determining gene, *Mdmd*, and we therefore defined this scaffold as corresponding to the Y^M^ chromosome, rather than the X chromosome (Figure 1).

To evaluate the completeness of the assembled genomes, we employed Benchmarking Universal Single Copy Orthologs (BUSCO) with a dipteran lineage dataset. All five strains had >97.5% complete BUSCO scores, suggesting that the assemblies contain nearly complete genomes. The MY strain genome, which was built from sequences of nine males (of the same strain), had a higher duplicated BUSCO score than the genomes of the other four strains, which were generated from single males. Using data from nine MY males probably caused incomplete filtering of the haplotype contigs in the *Purge_dup* program, compared to the other strains, which increased the duplication rate in the final genome assembly of the MY strain. As BUSCO provides annotations only for lineage-specific core genes, we also ran EGAPx (Eukaryotic Genome Annotation Pipeline-External) in order to generate complete gene annotations of each of the five strain assemblies (see details in Methods).

### Synteny conservation across schizophoran flies

We investigated synteny conservation of the house fly chromosomes with other schizophoran flies by comparing with *Drosophila melanogaster* and *Lucilia cuprina*. We first used a GENESPACE pipeline (see details in Methods), which tracks blocks of orthologous genes between assembled genomes. We assigned each of the five major house fly chromosomes (I-V) in our assemblies to Muller elements A-E based on the orthologous genes within synteny blocks with the other fly species [19–21] (Figure 2A, Figure S2A, B). Although we could find conserved syntenic blocks on element F between *D. melanogaster* and *L. cuprina*, we could not identify homology of element F with the house fly X chromosome (Figure 2A). The lack of homology between element F and the house fly X chromosome is consistent with prior work that failed to detect homology between element F and the X in Muscidae [30,31]. However, our GENESPACE analysis relied on homology between blocks of genes, and it may have missed gene-level homology in smaller segments of the chromosome.

**Figure 2.**
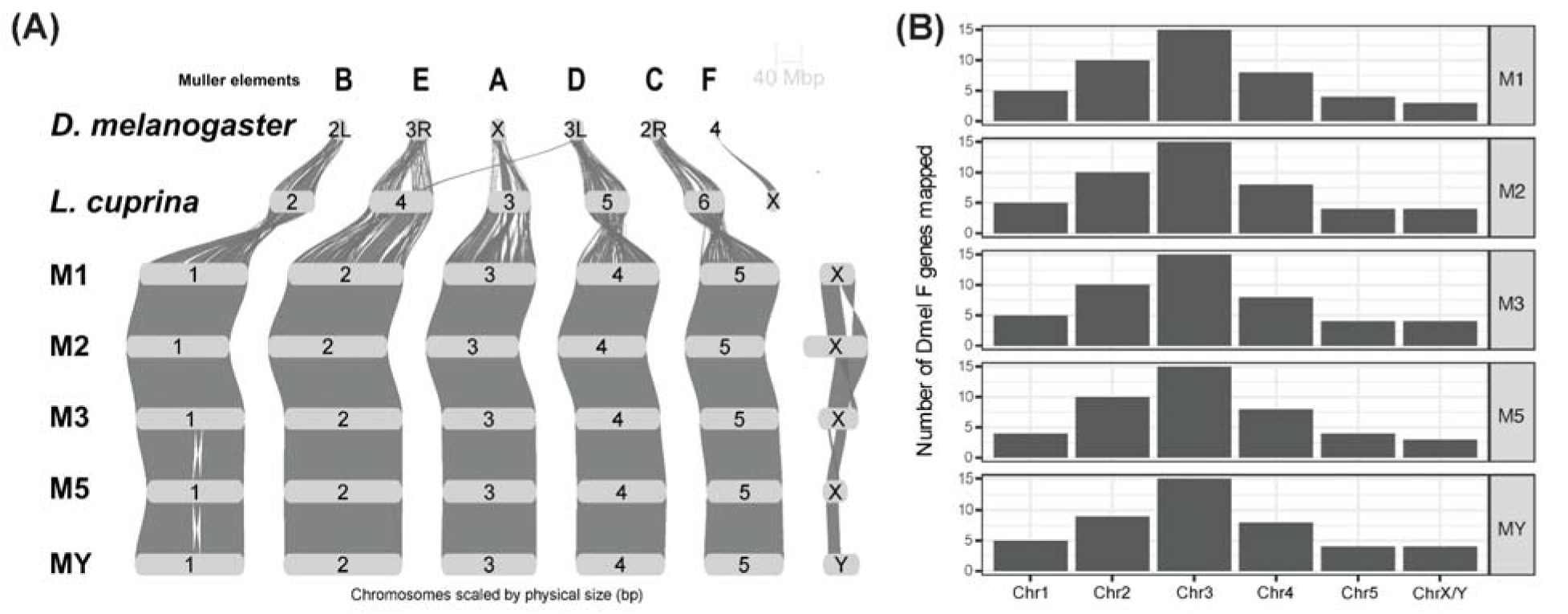
(A) Riparian plot showing synteny blocks among *D. melanogaster*, *L. cuprina*, and house fly (*M. domestica*) strains. (B) Assignment of *D. melanogaster* Muller element F gene to chromosomes of the five house fly strains by reciprocal best blast hits.

To address the possible shortcomings of the GENESPACE approach, we next identified individual homologous genes between the house fly X chromosome and Muller element F. To this end, we employed BLASTP to search for homologs between the house fly, *D. melanogaster*, and *L. cuprina* protein-coding genes. For house fly chromosomes I–V, the vast majority of genes had a best hit on the homologous Muller element in the *D. melanogaster* and *L. cuprina* genomes (Figure S2A, B), demonstrating homology of Muller elements A–E across the three species. In contrast, genes annotated on the house fly X chromosome had best hits across all six *D. melanogaster* and *L. cuprina* elements (Figure S2C,D), consistent with the lack of homology between the house fly X chromosome and element F.

We next assessed if we could find any remnant of shared ancestry between the house fly X chromosome and the *L. cuprina* X chromosome. Our approach calculated the proportion of genes on a chromosome in one species that had their best blast hit on the hypothesized homologous chromosome in the other species. Applying this method to *L. cuprina* and *D. melanogaster,* we found that 54-60% of *L. cuprina* X chromosome genes had their best hit on *D. melanogaster* element F, or *vice versa* (Figure 3A, B), demonstrating homology of the two chromosomes. In contrast, of all the *L. cuprina* X-linked genes with a blast hit in the house fly genome, 7-10% of the *L*. *cuprina* X chromosome genes had a best hit on the house fly X chromosome across all of our assemblies (Figure 3C-F). This is much smaller than the fraction of element F genes shared between *L. cuprina* and *D. melanogaster*. However, the *L. cuprina* and house fly X chromosomes have only 1.5-2% of all genes in the genome. If we use the proportion of genes on the X chromosomes as a null expectation, 7-10% best hits between the house fly and *L. cuprina* X chromosomes is a significant excess (*X*^2^_1_ > 34, p<10^-8^, Table S1). This provides evidence for homology between element F and the house fly X chromosome, although much less conservation than between *L. cuprina* and *D. melanogaster*.

**Figure 3.**
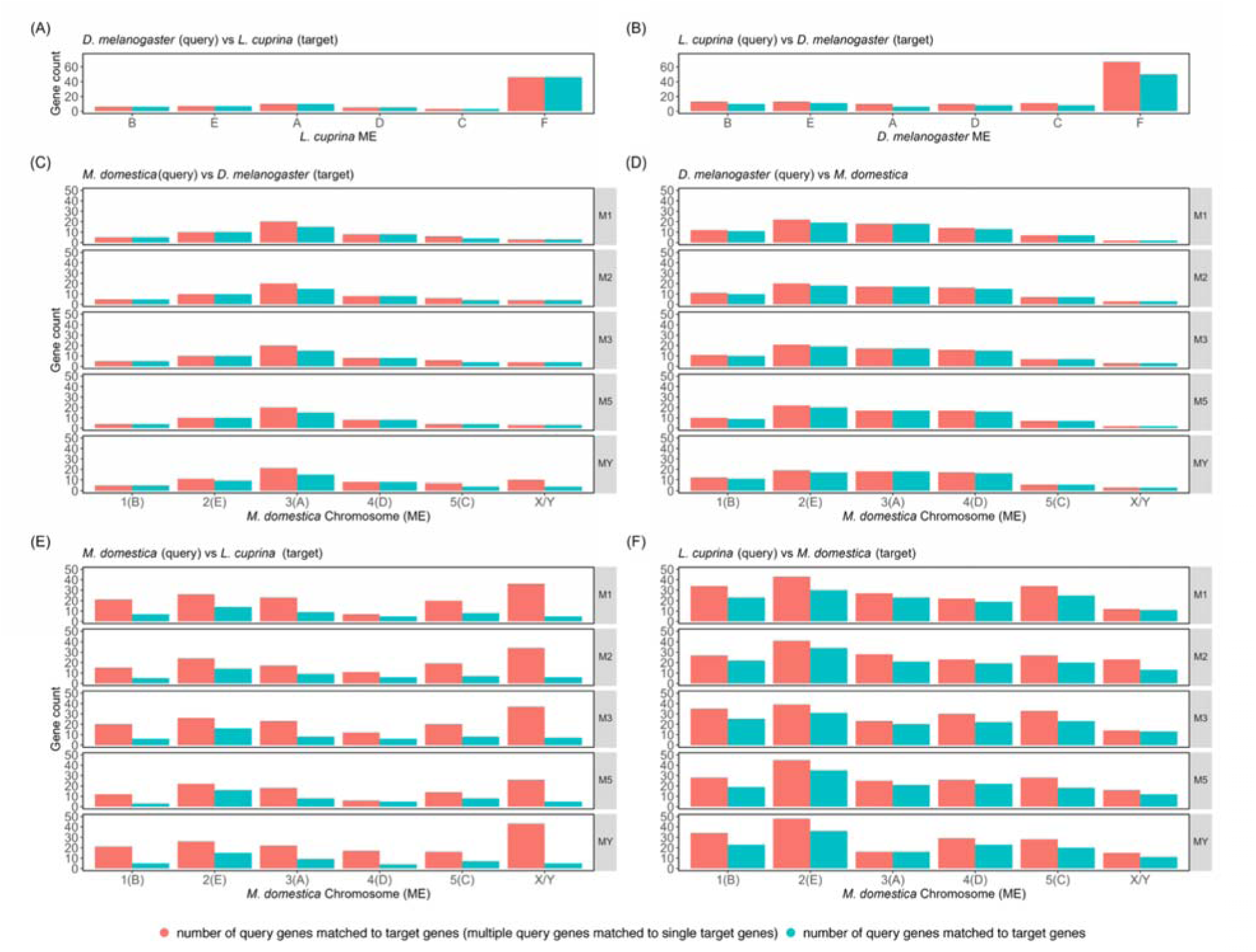
Chromosomal distribution of the Muller element (ME) F genes by BLASTP matches between two species. (A) Best blast hits of *D. melanogaster* genes to *L. cuprina* genes, and then selection by *D. melanogaster* F genes. (B) Best blast this of *L. cuprina* genes to *D. melanogaster* genes, and then selection by *L. cuprina* F genes. (C) Best blast hits of the house fly genes to *D. melanogaster* genes, and then selection by *D. melanogaster* F genes. (D) Best blast hits of *D. melanogaster* genes to the house fly genes, and then selection by *D. melanogaster* F genes. (E) Best blast hits of the house fly genes to *L. cuprina* genes, and then selection by *L. cuprina* F genes. (F) Best blast hits of *L. cuprina* genes to the house fly (*M. domestica*) genes, and then selection by *L. cuprina* F genes.

### Variation across X chromosomes and differentiation from Y^M^

The poor conservation of the house fly X chromosome with element F led us to further investigate the house fly X and Y^M^ assemblies. Previous cytogenetic studies demonstrated that the house fly X chromosome is comparable in length to the second chromosome [40,41], which is the largest chromosome in the house fly genome [42]. However, the X chromosomes in our genome assemblies were shorter than the other five chromosomes (Table 1), suggesting that the X assemblies might be incomplete. This is probably because the house fly X chromosome is enriched for repetitive DNA sequences [40,41], which can be difficult to sequence and assemble [43].

We tested if the incomplete X or Y^M^ chromosome assemblies could explain the variation in assembled genome sizes among strains. The assembled genomes of the strains differed by >100 Mb (Table 1), which is 12.4% of the average size of the assembled genomes. We compared the coefficient of variation (CV) of chromosome sizes across strains with the entire genome (CV=5.03) to evaluate the extent of variation in the assembled size of each chromosome. The assemblies of chromosomes II–V varied much less than the genome as a whole (0.9 < CV < 2.5), while chromosome I assemblies had a similar amount of variation as the total genome (CV=5.27). In contrast, the assembled X or Y^M^ chromosomes varied by >45 Mb (Table 1), or 63% of the average length of each chromosome (CV = 24.0). This amount of variation across X and Y^M^ chromosome assemblies is half of the total variation in assembled genome sizes across strains, despite the X or Y^M^ chromosomes representing <20% of the total genome size. Therefore, the X or Y^M^ chromosomes were outliers in the variation of their assembled sizes, which could explain much of the variation in total assembled genome sizes.

We evaluated if the variation in assembled X chromosome sizes could be associated with differences in the sequence content and organization of the X across strains. Chromosomes I–V were almost perfectly collinear across strains, with occasional inversions differentiating strains or haplotypes (Figure 2A, and see below). In contrast, multiple inversions and other rearrangements differentiated the X and Y^M^ chromosomes across strains, which are detectable using either gene-wise or sequence-based alignments (Figure 2A; Figure S3). However, many of the X chromosome inversion breakpoints are located at junctions between contigs (Figure S3), suggesting that they may be assembly artifacts. Nonetheless, one aspect of the X chromosome variation cannot be explained by assembly errors: the X chromosome in the M2 strain has a region not present in the other strains (Figure 2A), which appears to be an expanded repeat that has lengthened the left arm (Figure S3).

To further evaluate the causes of variation across assembled X and Y^M^ chromosomes, we estimated the amount of sequence data generated and used for our chromosome assemblies. We mapped PacBio HiFi reads from males of each strain to the genome assembly of the corresponding strain to quantify relative sequencing coverage for each chromosome. We normalized the sequencing coverage of a chromosome in each strain by the median coverage of the five (I-V) chromosomes of the same strain to compare the coverage along the chromosomes among strains (see details in Methods). Expected normalized coverage is 1 if all sites were equally sampled, and a normalized coverage value of 0.5 indicates that a chromosome was present at half the copy number as the others (i.e., haploid instead of diploid). In the M2, M3, and M5 strains, the median coverage of the X chromosome was below 1, but above 0.5; each median is 0.718, 0.843, and 0.802 (Figure 4A, Figure S4). These coverage values less than 1 on the M2, M3, and M5 X chromosomes can probably be explained by read mapping issues. The enrichment of repetitive sequences on the X chromosome [40,41] likely leads to mapping ambiguity, where sequencing reads map to multiple genomic locations.

**Figure 4.**
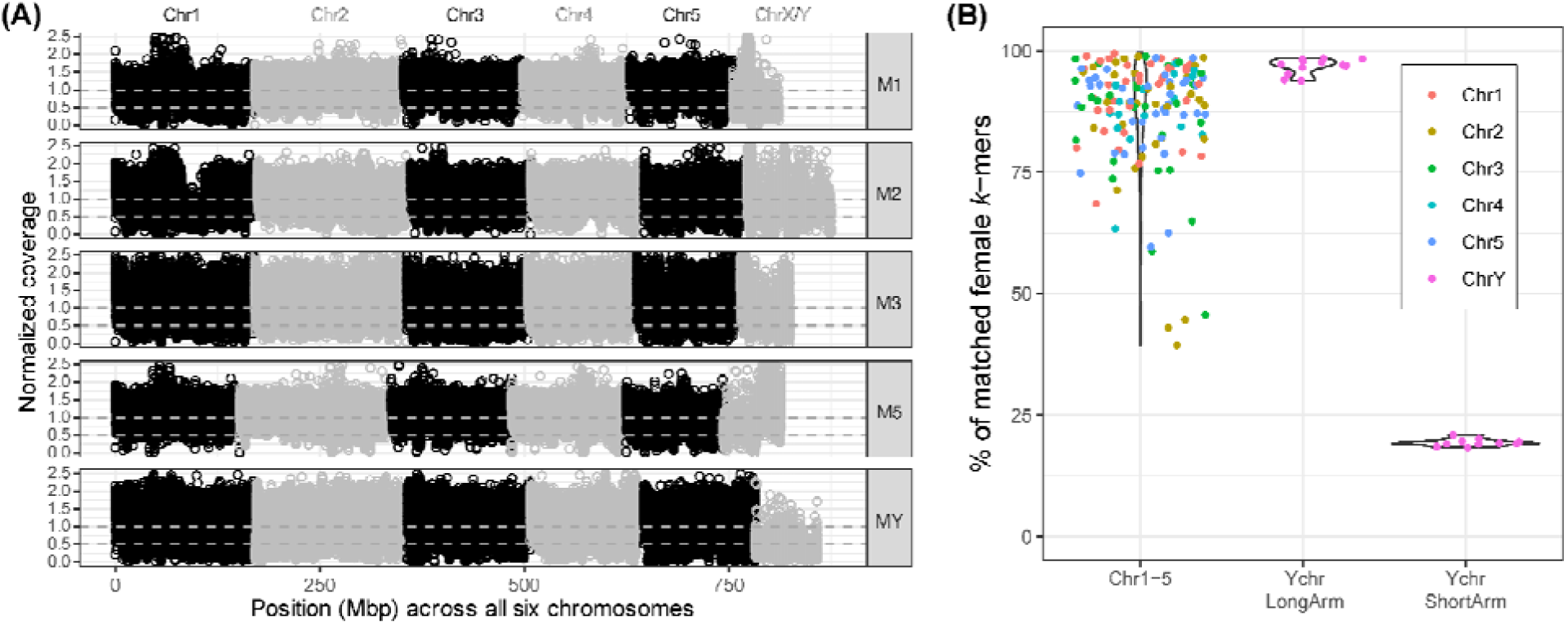
(A) Normalized coverage of male reads to the chromosomes in the genome assembly of the strain from which the reads were sampled. The alternating black and grey represent each chromosome. For the MY strain, the sixth chromosome is Y^M^. (B) Percentage of female read (15-mer) matches with male genome assembly in the MY strain. Points represent each contig of the MY male assembly and colors chromosomes (scaffolds corresponding to chromosomes).

Ambiguously mapped reads were discarded in the sequencing coverage analysis, which could have led to an underestimate of mapping coverage on the X chromosome. When considering the issues, these results indicate that the M2, M3, and M5 X chromosomes were likely sequenced to equivalent depth as the autosomes.

Unlike the M2, M3, and M5 strains, the median coverage of the X or Y^M^ chromosome in the M1 and the MY strain were below 0.5. The coverage of the X chromosome in the M1 strain was 0.434, and the coverage of the Y^M^ chromosome in the MY strain was 0.412 (Figure 4A, Figure S4). The median coverage <0.5 is consistent with a single copy of the Y^M^ chromosome in the MY strain (Figure 4A, S4), which we investigated further (see below). The low coverage of the M1 X chromosome could be explained by a reduced X copy number if the sampled M1 male had only one copy of the X chromosome. This could be possible because XO house flies are viable and fertile [41,44]. The M1 strain had the smallest assembled X chromosome of any strain (Table 1), consistent with sampling less X chromosome DNA from an XO male and, therefore, a reduced assembly size. In addition, we were unable to generate a haplotype phased assembly for the X chromosome from the M1 male, which would occur if the individual had an XO genotype. However, we also failed to generate haplotype phased assemblies of the X chromosomes in the M2, M3, and M5 strains, where there was X-coverage closer to the diploid expectation. Additional sequencing and cytogenetic study is thus required to determine if the X chromosome in the M1 strain is haploid or abnormal.

We also tested for sequence differences between the X and Y^M^ chromosomes, which could explain our coverage results above. Cytogenetic studies identified conspicuous morphological differences between the house fly X and Y^M^ chromosomes [35,40,41]; however, no differences between the X and Y^M^ chromosomes were observed in a short-read sequencing study using coverage and *k*-mer approaches [36]. To investigate this discrepancy, we analyzed our long-read sequencing and high-quality assemblies in combination with publicly available whole-genome sequencing data of males and females from the same MY strain [36]. The house fly Y^M^ chromosome has two arms, one short and one long, defined by a distinguishable C-banding pattern [41]. *Mdmd* is located in a pericentric region of the Y^M^ short arm, which has distinct C-banding from the long arm [35,41]. We used the contact map of the Y^M^ chromosome in the MY strain to identify the chromosome arms and centromere, and we used the location of *Mdmd* to define the long and short arms (Figure 1, Figure S5G: red arrow). In accordance with the previous studies, we found *Mdmd* near the centromere of the Y^M^ chromosome (Figure S5G: red arrow).

We next used a *k-*mer comparison approach and gene content analyses to identify Y^M^-specific sequences [45]. We first tested if contigs derived from the Y^M^ chromosome contained sequences enriched in males relative to females (Table S2). The long arm of the Y^M^ chromosome was highly matched by female sequences (>90% *k*-mer matches), resembling an autosome. In contrast, a small percentage (∼20%) of the Y^M^ short arm was matched by female *k*-mers, suggesting enrichment of male-specific sequences in the Y^M^ short arm (Figure 4B). We annotated 197 genes on the long arm of the Y^M^ chromosome and 92 genes on the short arm.

We asked if these are similar in sequence to genes found on the X chromosomes of other strains. Using a 95% sequence identity cutoff, we found that 45-63% of genes on the long arm of the Y^M^ chromosome have a similar gene on the X chromosome of another strain, while only 3-12% of genes on the short arm have an X chromosome match (Table 2). These results indicate that the long arm of the Y^M^ chromosome is homologous to the X chromosome. The Y^M^ short arm, where *Mdmd* is located, is likely specific to the MY males, which is consistent with the previous cytological work [35,41].

**Table 2.** Y^M^ genes with >95% sequence identity to X chromosome genes of other strains.

| strain | Long arm<br>(197 genes) | Short arm<br>(92 genes) |
| --- | --- | --- |
| M1 | 121 (61%) | 3 (3%) |
| M2 | 100 (51%) | 6 (7%) |
| M3 | 124 (63%) | 11 (12%) |
| M5 | 88 (45%) | 5 (5%) |

### Polymorphic chromosomal inversions

We identified paracentric inversions in three strains that differentiated the gene order within chromosomes I-V. A 12.4 Mbp inversion on chromosome I was homozygous in the M5 strain (Figure 2A, Figure S6), and the same chromosome I inversion was heterozygous in the M2 strain (Figure S6). We also identified a 35.7 Mbp heterozygous inversion on one arm of chromosome II in the M2 strain (Figure 5A). Lastly, a 38.4 Mbp inversion was observed on one arm of chromosome V in both the M2 and MY strains. At least one of the inversion breakpoints on chromosomes I, II, and V were identified in the middle of assembled sequences (within contigs), not in assembly gaps between scaffolds (Figure 5; Figure S6), suggesting that the inversions were not artifacts of scaffolding.

**Figure 5.**
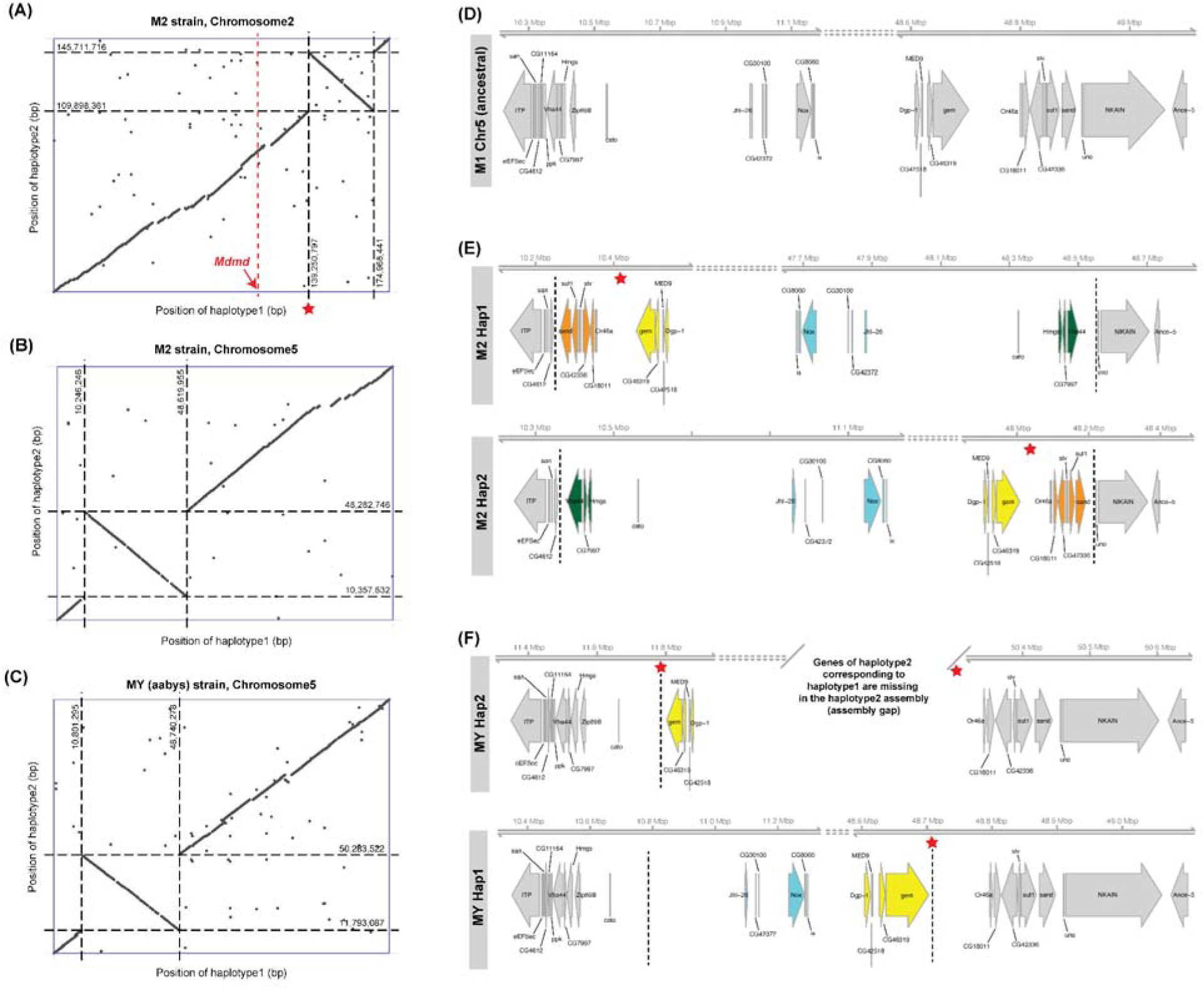
Polymorphic inversions in chromosomes II and V of the M2 (A, B) and in chromosome V of MY (C) house fly strains. Synteny in chromosome V inversion breakpoints from the M1 (D), M2 (E) and MY (F) strains; *D. melanogaster* gene symbols, homologous to the house fly genes, were used. Red stars represent the locations of assembly gaps on scaffolds.

Only the chromosome II inversion in the M2 strain was on a proto-sex chromosome. The remaining inversions were on autosomes in each strain from which they were sampled. Inversions on Y chromosomes are hypothesized to maintain genetic linkage between a male-determining locus a male-benefiical sexually antagonistic allele [7]. However, the chromosome II inversion in M2 does not capture *Mdmd*, which is located in the pericentromeric region, approximately 30.3 Mbp proximal to the closest inversion breakpoint (Figure 5A).

Both the M2 and MY strains are heterozygous for an inversion on chromosome V, capturing very similar genomic regions (Figure 5B, C). This suggests that the strains share the same inversion on chromosome V. However, the breakpoints in the M2 strain fall between different genes than the breakpoints in the MY strain (Figure 5D-F). We therefore conclude that the M2 and MY strain carry independently derived inversions that happen to capture very similar regions of chromosome V. In contrast, the breakpoints of the chromosome I inversion fall between the same genes in the M2 and the M5 strains (Figure S6), suggesting that the inversion on chromosome I in the M2 and M5 strains has a single origin.

### Differentiation of proto-X and proto-Y chromosomes

We further explored divergence of the I^M^, II^M^, III^M^, and V^M^ proto-Y chromosomes from their corresponding proto-X chromosomes by measuring sequencing coverage across the genome. We excluded the Y^M^ chromosome from this analysis because of the large-scale structural variation described above. As expected for autosomes, the median coverages across chromosomes I-V for most strains were close to the expected value of 1 for a diploid chromosome (Figure 4A). Proto-X and proto-Y chromosomes are expected to have similar coverage as autosomes because they are largely undifferentiated from each other aside from a male-determining locus [46]. However, the median coverages of the proto-sex chromosomes in the M1, M3, and M5 strains were slightly below one (Figure 4A, Figure S4, medians are 0.809, 0.876, and 0.896, respectively), suggesting moderate yet incomplete X-Y divergence.

Alternatively, there may be read mapping issues on the proto-Y chromosomes that are associated with allelic divergence with their proto-X chromosomes (see below), which may interfere with the accurate mapping of reads [32].

Young proto-sex chromosomes are also expected to have elevated heterozygosity as a result of allelic differences between the proto-X and proto-Y [19,46]. To measure heterozygosity, we calculated the pairwise differences between haplotypes within each strain in sliding windows along chromosomes I-V, sampling only synonymous sites within protein-coding genes (Figure 6, Table S3). We observed similar patterns when we sampled heterozygosity across the entire chromosomes in both the entire protein-coding DNA regions and all non-coding sites (Figure S7, Table S4). The heterozygosities within the M1, M2, M3, and M5 strains were compared to that of the MY strain as the baseline reference for each protein-coding gene because chromosomes I-V are autosomal in the MY strain [36]. Similar results were obtained without using MY as a reference (Figure S7), although we note a few differences below. We observed elevated heterozygosity across much of the proto-sex chromosome in the M1 and M3 strains (Figure 6), which is evidence for X-Y divergence in each strain. In contrast, we only observed elevated heterozygosity in the M2 strain within the region containing the polymorphic chromosome II inversion (Figure 6, Figure S7). In addition, elevated heterozygosity in the M5 strain was limited to only one arm of chromosome V, likely because we used the MY strain as a reference (Figure 6). The MY strain had elevated heterozygosity within the inverted region of chromosome V (Figure S7, S8), which increased the baseline level to which the M5 strain was compared. When we compared heterozygosity across chromosomes in M5 males without using MY as a reference, we indeed observed elevated heterozygosity across the entire length of chromosome V (Figure S7). These results show that X-Y divergence has occurred across the entire proto-sex chromosomes in most strains. The one exception is the M2 strain, where the X-Y divergence was confined to one arm of the second chromosome where there is a heterozygous inversion.

**Figure 6.**
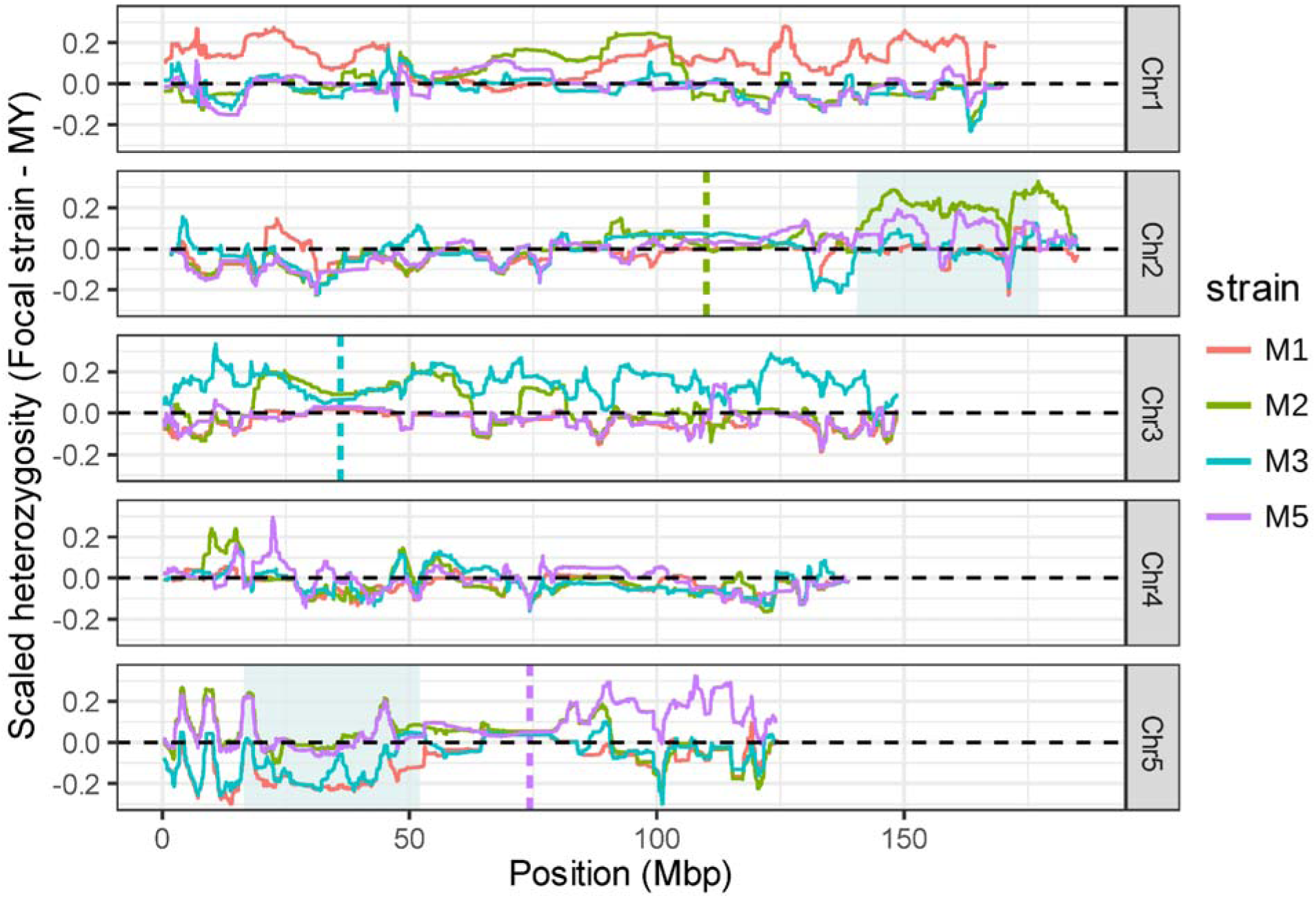
Synonymous divergence (*K_s_*) between the homologous chromosomes of the house fly strains. The chromosomal divergence of M1, M2, M3, and M5 strains was displayed in reference to the MY strain (see Methods for details). Cyan shaded regions indicate heterozygous inversions in chromosome 2 and 5 of the M2 strain.

Another feature of X-Y divergence is the accumulation of repetitive sequences on the Y chromosome [7]. Thus, we examined whether the house fly proto-Y chromosomes acquired more repetitive sequences compared to their homologous proto-X chromosomes. We found that chromosomes I-V are composed of ∼70% repetitive DNA sequences (Figure S9), consistent with prior observations that more than half of the house fly genome consists of interspersed repeats [47]. However, repeat composition, including transposable elements and satellites, was similar between proto-Y and proto-X chromosome haplotypes across all the strains (Figure S9A).

This analysis is somewhat limited by uncertainties in phasing the X and Y haplotypes, and we therefore also compared proto-sex chromosomes with the euchromatic autosomes within each strain. Within each strain, the repeat composition in the proto-sex chromosomes was not conspicuously different from that in autosomes of the same strain (Figure S9A). These results suggest that there is not a disproportionate accumulation of repetitive DNA on the house fly proto-Y chromosomes.

### Evolution of *Mdmd* and *M* loci

The male determining locus is the earliest evolving and sometimes only feature of a young proto-Y chromosome that differentiates it from the proto-X [48]. Therefore, to determine a maximum age of the house fly proto-Y chromosomes, we aimed to refine the age of the house fly male determining gene, *Mdmd*. Previous work found that *Mdmd* is a paralog of spliceosomal factor gene CWC22, also known as *nucampholin* (*ncm*) in *D. melanogaster* [33]. The duplication of *ncm* occurred after the divergence of *M. domestica* and *Stomoxys calcitrans* [33], but it was not clear when *Mdmd* arose in the ∼35 My since the *Musca* and *Stomoxys* genera diverged. To refine the date of the duplication event, we generated a phylogenetic tree with *ncm* genes from fly species within the family Muscidae (including other *Musca* species), along with *Mdmd* and *ncm* of each house fly strain. The phylogenetic tree indicates that the *Mdmd* duplication occurred after divergence of *M. domestica* from *Musca vetustissima* (Figure 7A).

**Figure 7.**
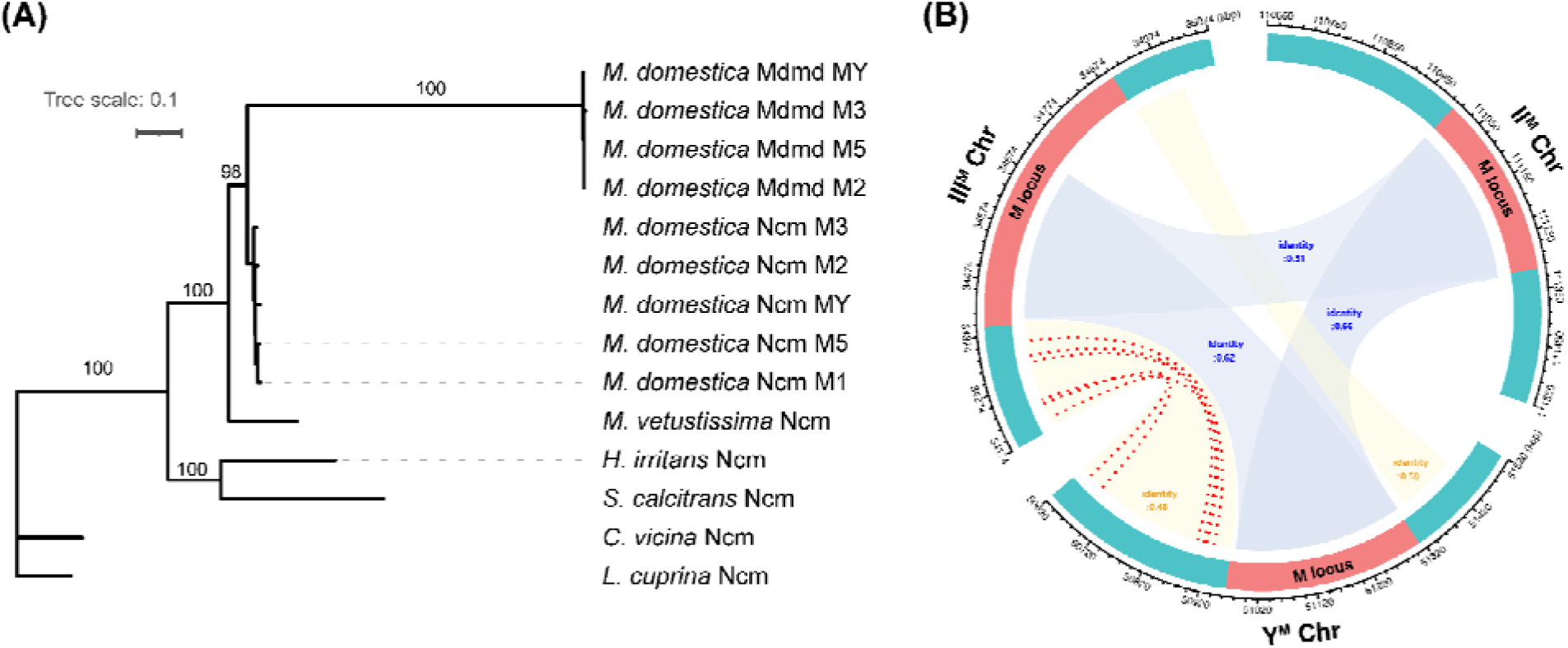
(A) Phylogenetic tree of Mdmd and Ncm proteins in calyptratae flies. Bootstrap values >85 are displayed at the corresponding nodes. (B) *M* locus and its flanking (*M*-adjacent) regions in the II^M^, III^M^, and Y^M^ chromosomes. Matched regions of *M* locus and *M*-adjacent sequences between proto-Y chromosomes (II^M^ vs Y^M^ or III^M^ vs Y^M^ or II^M^ vs III^M^) are light blue-shaded and light yellow-shaded. Identity represents sequence similarity between the proto-Y chromosomes. The coding sequence matches are represented as red dotted lines with >0.99 identity.

*M. ventustissima* belongs to the *Musca sorbens* species complex, which is estimated to have diverged from *M. domestica* 15.6 My ago (Mya) [49–51]. This suggests the duplication of house fly *Mdmd* occurred less than 15 Mya, and therefore the house fly proto-Y chromosomes are likely less than 15 My old.

Previous work also identified tremendous variation in the structure of the *M* loci across the II^M^, III^M^, V^M^, and Y^M^ chromosomes [35]. However, that analysis lacked clear resolution of the *M* loci and flanking regions because it relied upon more fragmented, contig-level assemblies than ours. To better understand the structural similarities and differences between the *M* loci, we examined *M*^II^*, M*^III^, and *M*^Y^. We selected these three loci because they have a similar composition of a single intact *Mdmd* and multiple truncated copies of *Mdmd* [33,35]. We excluded the *M*^V^ locus from our comparisons because it has a completely different organization from *M*^II^, *M*^III^, and *M^Y^* [35]. In our analysis, we defined the *M* locus as the sequence spanning all *Mdmd* copies (both complete and partial), and we refer to sequences on either side of the *M* loci as *M-*adjacent.

We discovered differences in the structure of *M-*adjacent regions that were not previously identified using the less contiguous genome assemblies. First, we confirmed a prior finding [35] that the *M*^III^-locus has an expanded region that is not found in the *M*^II^ or *M*^Y^ loci (Figure 7B). We also discovered that the III^M^ and Y^M^ chromosomes share *M-*adjacent sequences that are not present in the II^M^ chromosome (Figure 7B). We found six protein coding genes in the *M*- adjacent sequences of the III^M^ and Y^M^ chromosomes, which have a sequence identity of >0.99 between the two strains. These identities are similar to the coding sequence divergence between the two strains on other chromosomes (Figure 7B, Table S5). We ran Interproscan (https://www.ebi.ac.uk/interpro/) with the peptide sequences of the six genes [52], and we found that five encode Ribose-phosphate pyrophosphokinase proteins. This enzyme converts ribose 5-phosphate into phosphoribosyl pyrophosphate, which is involved in purine synthesis [53,54].

## Discussion

Recent studies across diverse taxa revealed multiple non-canonical trajectories of sex chromosome evolution [18,55,56]. To test hypotheses about canonical and non-canonical evolutionary trajectories within a single genomic environment, we sequenced and assembled five different house fly genomes, each of which carried a different proto-Y chromosome.

Through phylogenetic analysis of the *Mdmd* gene and its ancestral paralog, we dated the origin of the house fly proto-sex chromosomes to less than 15 Mya. We also confirmed that the house fly X chromosome is highly divergent from Muller element F, and the corresponding Y^M^ chromosome has a Y-specific arm that is not homologous to the X chromosome. We identified one inversion on a proto-Y chromosome, but most of the proto-sex chromosomes lacked any large inversions. Despite the lack of inversions, we detected X-Y divergence across the entire lengths of most of the proto-sex chromosomes.

### Evolution of *M* loci

We were able to assemble both the *M* loci and *M-*adjacent sequences in each strain, allowing us to gain new insights into the origins of sex determining loci. Previous work demonstrated that the *M*^III^ locus is larger than *M*^II^ and *M*^Y^ because *M*^III^ has more repetitive sequences flanking the functional copy of *Mdmd* [35]. Many of the repeats at each *M* locus are pseudogenized duplications of *Mdmd*, and there is only one functional copy of *Mdmd* in each *M*^II^, *M*^III^, and *M*^Y^ locus [35]. We confirmed that the *M*^III^ locus has an expanded region that is not found in the *M*^II^ or *M*^Y^ loci. Using assemblies of each locus, we also found that the sequence identity between *M*^Y^ and either *M*^II^ or *M*^III^ is greater than the identity between *M*^II^ and *M*^III^ (Figure 7B). In addition, we discovered that there are *M-*adjacent sequences shared between the III^M^ and Y^M^ chromosomes that are absent in the II^M^ chromosome (Figure 7B). The shared sequences across *M* loci and *M-*adjacent regions suggest that the loci on each chromosome arose from the duplication of large segments from one proto-Y chromosome to the other.

Our comparisons of the contents of the *M* loci and *M-*adjacent regions are suggestive of the evolutionary order in which the proto-Y chromosomes arose. The shared *M-*adjacent sequences between Y^M^ and III^M^ chromosomes suggest that the *M* locus on one chromosome arose from the other by a large segmental duplication, but the ancestral locus is ambiguous. Additionally, *M*^Y^ and *M*^II^ have the highest sequence identity, suggesting that one of these is the most recently derived *M* locus out of the three under discussion here. We hypothesize that *M*^II^ arose from *M^Y^* because the Y^M^ chromosome must have existed earlier to participate in the duplication that created the shared *M*-adjacent sequences between Y^M^ and III^M^. However, we are unable to determine the order of duplications that created *M*^Y^ and *M*^III^.

Our results demonstrate how new sex chromosomes originated within the house fly genome via segmental translocations encompassing both large and small chromosomal segments. Prior work established that the *M^V^* locus contains two functional copies of *Mdmd* in an inverted tandem orientation [35], suggesting that a duplication event at the scale of a single gene (<10 kb) gave rise to M^V^. In contrast, the shared sequences between M^Y^ and M^II^ are consistent with a 300 kb segmental duplication that captured a large portion of an existing *M* locus.

Moreover, the alignment of both the *M* locus and flanking M-adjacent sequences between M^Y^ and M^III^ provide evidence that a segmental duplication >800 kb is responsible for the creation of one of those *M* loci. Lastly, the lack of an *Mdmd* gene on the I^M^ chromosome suggests an entirely different process is responsible for the creation of the *M*^I^ locus. These findings demonstrate that there is no single canonical process to generate a sex determining locus, even within a genome that re-uses the same gene as a male-determining factor.

### Evolution and origins of the house fly X and Y^M^ chromosomes

The X and Y^M^ chromosomes are the most non-canonical sex chromosomes in the house fly genome. While many fly taxa have an X chromosome that is homologous to Muller element F [19,22], recent studies failed to identify homology between the Muscidae X chromosome and element F [30,31]. We similarly found very little evidence that the house fly X (or Y^M^) chromosome is homologous to element F. For example, although many of the house fly X/Y^M^ genes had identifiable homologs in the *D. melanogaster* or *L. cuprina* genomes, very few of those homologs were on element F (Figure 2B, S2A,B). This suggests that the X and Y^M^ chromosomes are either highly diverged from Muller element F or arose independently of element F. An independent origin of the house fly X chromosome could have involved a B chromosome acquiring a male-determining locus [57], similar to how polymorphic B chromosomes carry sex-determining genes in cichlid fish [58,59].

As a test for element F ancestry of the house fly X chromosome, we compared our X and Y^M^ assemblies directly with the *L. cuprina* X chromosome. We selected this comparison for three reasons. First, element F is still an X chromosome in *L. cuprina*, but it has become an autosome in *D. melanogaster* [19]. Comparing house fly with *L. cuprina* therefore allows us to compare two X chromosomes. Second, *L. cuprina* is more closely related to house fly than *Drosophila*, and therefore a better reference point for comparing a highly divergent chromosome. Third, the *L. cuprina* X chromosome has more than twice as many genes as *D. melanogaster* element F, suggesting that element F has lost genes in the lineage leading to *Drosophila* [31]. The *L. cuprina* X chromosome is thus a more intact representation of element F. We did indeed identify a significant excess of *L. cuprina* X chromosome genes with best hits on the house fly X chromosome, consistent with the hypothesis that the house fly X is a highly divergent homolog of element F.

In addition to being divergent from element F, we also identified a lot of intra-specific variation of the house fly X chromosome. Across the four strains that are homozygous for the X chromosome (i.e., all but the MY males), multiple inversions appeared to differentiate the X chromosomes (Figure S3). Although many of these inversions may be assembly artifacts, we also identified expansions of large repetitive sequences, including one that makes up approximately half of the M2 X chromosome (Figure 2A, S3). The structural variation on the X chromosome appears to be far greater than on the other five chromosomes.

The M1 strain may have an especially peculiar X chromosome. The M1 X chromosome had sequencing coverage that was half of what is expected if it were diploid (Figure 4A; Figure S4). The I^M^ chromosome is also the only proto-Y that lacks *Mdmd* [33], suggesting that a different gene acts as the male-determining locus in that strain. We were unable to identify I^M^-specific sequences that could be candidate male-determining loci. It remains to be determined if the reduced sequencing coverage of the X chromosome in the M1 strain is associated with its peculiar male-determining locus or hemizygosity.

We lastly identified similarities and differences between the Y^M^ and X chromosomes that could help us understand the evolutionary origins of Y^M^. Synteny comparison with the X chromosomes of other strains (Figure 2A) and *k*-mer comparison between males and females in the MY strain (Figure 4B) point to the conservation of gene content and sequences in the long arm of the Y^M^ chromosome with the X chromosome. In contrast, the short arm of the Y^M^ chromosome appears to be present only in the MY males, suggesting that the short arm is Y^M^-specific. This is consistent with previous cytological work [35,40,41]. However, it is not clear whether the Y^M^- specific region was acquired along with the *M* locus as a fusion between two chromosomal segments, whether the Y-specific short-arm pre-existed the origin of *Mdmd,* or whether it arose after *Mdmd* as an outcome of X-Y divergence.

### Chromosomal inversions, X-Y divergence, and suppressed recombination

The I^M^, II^M^, III^M^ and V^M^ chromosomes appear to follow more canonical modes of sex chromosome evolution, albeit with some deviations. Multiple models of sex chromosome evolution predict inversions on Y and W chromosomes [18,60,61]. While we identified four polymorphic chromosomal inversions segregating in the strains we sampled, only one inversion was located on a proto-sex chromosome (Figure 5A). The one sex chromosome inversion was observed in the M2 strain, and there was elevated heterozygosity within and adjacent to the inversion on chromosome II (Figure 6, Figure S7). Allelic divergence is expected to arise as a result of suppressed recombination between proto-X and proto-Y chromosomes in males, and chromosomal inversions are hypothesized to be an important mechanism that evolves to suppress X-Y recombination [6,62,63]. Notably, chromosomal inversions are only expected to be required to suppress X-Y recombination if there is crossing-over during male meiosis, and, indeed, there is some evidence for male recombination in house flies with non-Y^M^ genotypes [64]. The II^M^ chromosome therefore appears to be evolving according to a canonical model for a Y chromosome.

Suppressed recombination via a chromosomal inversion is expected to be favored on a Y chromosome in order to maintain genetic linkage between the male-determining locus and male-beneficial alleles [4]. The chromosome II inversion is located near the *Mdmd* locus on the II^M^ chromosome, but *Mdmd* is proximal to (i.e., outside of) the inverted region. Nonetheless, recombination is also expected to be suppressed proximal to a paracentric inversion [65], suggesting that genetic linkage between *Mdmd* and alleles within the inversion may be elevated relative to the expectation based on their physical distance. The inversion on chromosome II, proximity to the male-determining locus, and elevated heterozygosity within the inverted region are all consistent with the expected relationships between chromosomal rearrangements, suppressed X-Y recombination, and X-Y divergence in a canonical sex chromosome.

The elevated heterozygosity on chromosome II is unlikely to be a technical artifact for the following reasons. We also observed elevated heterozygosity within autosomal inversions on chromosome I (in M2 males) and chromosome V (in M2 and MY males), similar to the chromosome II inversion in the M2 strain (Figure 5, 6, S7). The inversions on chromosomes I, II, and V are heterozygous in each strain where they are found, demonstrating divergence between inverted and standard arrangements. This is the expectation if recombination is suppressed and regardless of whether the inversions are autosomal or sex-linked linked [66,67]. In comparison, the M5 strain is homozygous for an inversion on chromosome I (Figure 2A), and heterozygosity is not elevated on chromosome I in the M5 strain (Figure 6, S7). We do not expect elevated heterozygosity when a strain is homozygous for an inversion, indicating that the observed elevated sequence heterozygosity in inversion heterozygotes is not an artifact of an inverted region on the chromosome.

One of the autosomal inversions we identified was shared across strains in a way that was consistent with the geographical origin of those strains. The M2 and M5 strains shared the same inversion on chromosome I. Both strains were collected in Pavia, Italy, suggesting that the chromosome I inversion segregates in this population. In contrast, the M2 and MY strains both have an inversion on chromosome 5, but the breakpoints differ between strains. The MY strain we sampled is known as aabys. It carries a recessive mutation on each of the euchromatic chromosomes (I-V), each of which has been described in laboratory cultures since the mid- 1950s. The different origins of the M2 strain (a wild population in Italy) and the MY strain (long-term laboratory culture) are consistent with those strains carrying independently derived chromosome 5 inversions.

We also observed elevated DNA sequence heterozygosity across the entire lengths of the proto-sex chromosomes in the M1, M3, and M5 strains (Figure 6, S7). Previous work identified elevated heterozygosity on the third chromosome in M3 males [36,68]. However, without a chromosome-scale assembly, the prior work lacked the resolution to determine if the pattern was chromosome-wide or restricted to specific segments. Our results demonstrate the III^M^ proto-Y chromosome is diverged from its homologous proto-X across its entire length, as are the I^M^ and V^M^ chromosomes (Figure 6, S7). Despite the elevated heterozygosity, we did not detect any inversions on the proto-sex chromosomes in the M1, M3, and M5 strains. Therefore, house fly proto-X and proto-Y chromosomes can diverge from each other without chromosomal inversions to suppress recombination.

X-Y divergence requires some form of suppressed recombination in XY heterozygotes. The lack of inversions on the I^M^, III^M^, or V^M^ chromosomes, along with the elevated heterozygosity on those chromosomes, suggests that another mechanism must suppress recombination. A likely candidate is low or no recombination in males carrying the I^M^, III^M^, or V^M^ chromosomes. Meiotic chromosomal segregation in males of many fly species, including members of the *Drosophila* genus, occurs via an achiasmate mechanism in which there is no crossing over [69]. Evidence for male recombination in house flies is mixed [70], with studies reporting male recombination rates ranging from zero to half the rate in females [64,71,72]. The elevated heterozygosity on the I^M^, III^M^, or V^M^ chromosomes is consistent with the zero estimate of the male recombination rate. However, as described above, the relationship between a chromosomal inversion and heterozygosity on the II^M^ chromosome is consistent with male recombination greater than zero. How can we reconcile these conflicting results?

We propose XX hypotheses for the conflicting associations between inversions and heterozygosity across house fly proto-sex chromosomes. First, male-recombination in house flies may be limited to II^M^ males. Second, even if there is no male recombination, the population-level recombination rates may differ across house fly proto-Y chromosomes because they can recombine with a proto-X when transmitted through females. This is because house fly females can carry a proto-Y chromosome if they also have the female-determining *Md-tra^D^* allele [70]. X-Y recombination may thus occur in females who carry both a proto-Y chromosome and *Md-tra^D^*, similar to the effects of female recombination in sex-reversed XY tree frogs [12–14]. If X-Y recombination occurs more often in *Md-tra^D^* females with a II^M^ chromosome, the population-level recombination rate of the chromosome II proto-X and proto-Y chromosomes would be higher than other proto-sex chromosome pairs. Nonetheless, a lower population-level recombination of all proto-Y chromosomes is expected if there is low or no male recombination and biased transmission of the proto-Y through males [73,74]. Additional experiments are required to test these hypotheses.

The location of *M* loci near the centromeres may further reduce the effects of X-Y recombination in females. Our chromosome-scale assemblies confirmed previous cytogenetic data that the *M* loci on chromosomes II, III, V, and Y^M^ are near the centromeres [35] (Figure S5).

Recombination tends to be lower near the centromere, relative to the interstitial regions of chromosome arms, in many animal genomes [75]. Therefore, the *M* loci may be inherently linked to the proto-Y chromosomes without any additional suppressors of recombination needed, even if they are occasionally transmitted through females [76]. However, we observed elevated heterozygosity along the entire lengths of the proto-sex chromosomes, which cannot be explained by proximity of *M* loci near centromeres. All of these factors—low or no male recombination, no inversions, centromeric proximity of *M*, and female transmission—mean that the house fly proto-sex chromosomes can appear to have canonical features of X-Y divergence via non-canonical means.

We refined the age of *Mdmd*, and therefore the house fly proto-Y chromosomes, to within 15 Mya (Figure 7A). As described above, we also observed minimal X-Y divergence of most of the house fly proto-sex chromosomes and no evidence for degeneration. In comparison, the *Drosophila miranda* neo-Y chromosome originated < 2 Mya, and there is extensive evidence that the neo-Y has experienced substantial degeneration [77,78]. Either the house fly proto-Y chromosomes are much younger than 15 My or they degenerate at a slower rate than *Drosophila* neo-Y chromosomes. The house fly proto-Y chromosomes may degenerate slower than *Drosophila* neo-Y chromosomes because the house fly proto-Y chromosomes can be transmitted through *Md-tra^D^* females. X-Y recombination in *Md-tra^D^* females (see above), along with male recombination, could allow for the purging of deleterious mutations from the proto-Y chromosomes, reversing the effects of Muller’s ratchet and hitchhiking [79,80]. This is consistent with a ‘fountain of youth’ model of sex chromosome evolution in which the differentiation and degeneration of a Y chromosome is prevented or mitigated by recombination with the X chromosome [81]. Testing if the house fly proto-Y chromosomes are young or degenerate slowly will require genomes of more closely related species and additional information about population-level recombination rates.

## Methods

### Housefly strains

We used five house fly strains (M1, M2, M3, M5, and MY) that each have a naturally occurring proto-Y chromosome. The locus carrying the male-determining gene, *Mdmd*, in the M2, M3, M5, and MY strains has been characterized in [33,35]. The M1 strain has an unknown male-determining factor (not *Mdmd*) on the first chromosome [33,34]. Males of each strain are heterozygous for their proto-Y chromosome: the M1 strain has I^M^/I males, the M2 strain has II^M^/II males, the M3 has III^M^/III males, the M5 strain has V^M^/V males, and the MY strain has Y^M^/X males. All strains have been maintained in laboratory settings for many generations and are highly inbred [35,82].

### PacBio HiFi Sequencing

We performed PacBio sequencing on the M1, M2, M3, and MY strains. For each strain, high molecular weight (HMW) DNA was extracted from a single male using the Qiagen MagAttract Kit (Qiagen, Hilden, Germany). A PacBio SMRTbell library was prepared separately for each male using the SMRTbell Express Template Prep Kit 2.0 (Pacific Biosciences, Menlo Park, CA, USA). The prepared library was bound and sequenced at the USDA Agricultural Research Service, Veterinary Pest Genetics Research Unit in Kerrville, Texas, on Pacific Biosciences 8M SMRT Cells with a Revio system (Pacific Biosciences, Menlo Park, CA, USA) (Table S6). See details in supplemental methods in additional file 1.

We used alternative approaches for the M5 and MY strains. First, we performed additional sequencing on the MY strain. HMW DNA was extracted from three individual males using the Monarch Extraction Kit (T3060) from New England Biolabs (NEB) (Table S6). The three separate HMW DNA samples were shipped to Novogene Corporation Inc., who prepared Pacbio sequel II DNA HiFi library for each DNA extraction and carried out the sequencing run on Pacbio sequel II (HiFi/CCS mode/cell). We found that we obtained better assemblies with these three samples than the previously described MY sequences or if we combined all MY sequence data. For that reason, we only used these three MY samples for our final assembly, but we deposited all PacBio data into the NCBI SRA database. We used publicly available PacBio HiFi sequencing data for the M5 strain males (Table S6) [35]. See details in supplemental methods in additional file 1.

### Hi-C sequencing

We generated Hi-C sequence data to scaffold the assemblies into chromosomes. For the M2 strain, a Hi-C library was generated from a single male by Arima Genomics and then sequenced on an Illumina NovaSeqXPlus using a high-throughput flow cell. For the remaining strains, a Hi-C library was generated from a single male by Phase Genomics using the Proximo Animal kit version 4.0. The MY sample was sequenced on a NovaSeq 6000 using an S4 flow cell. The M1, M3, and M5 samples were sequenced on a NovaSeqXPlus using a high-throughput flowcell. See details in supplemental methods in additional file 1.

Genome assembly.

For each strain, we first generated a draft genome by running *Hifiasm* (version 0.19.9) [83,84] with PacBio HiFi reads, which output each haplotype assembly and a primary assembly. For the primary assembly, we performed *Purge_dups* (version 1.2.5) [85] to remove haplotigs and then scaffolded the purged assembly using *YaHS* (version 1.2) [86] with the alignment of Hi-C reads generated through BWA [87], following Arima Genomics’ mapping pipeline (https://github.com/ArimaGenomics/mapping_pipeline). The resulting assemblies were visually reviewed and manually corrected for contig orders and orientations with *Juicebox* version 2.15 [88]. For diploid haplotype-phased assemblies, we applied the same procedures as for the primary assembly, except for the haplotigs removal step using *Purge_dups*. We first ran *Purge_dups* on the larger haplotype assembly of the two, and we next ran the same program on the second haplotype assembly with added sequences purged from the first haplotype assembly for more contiguous haploid genomes. We employed BUSCO version 5.6.1 to assess the completeness of five strain assemblies with the diptera_odb10 [89]. Assembly quality value (QV) and kmer completeness were calculated with 21-length kmers using Meryl v1.4 and Merqury v1.3 using the diploid option [90].

We generated Hi-C contact maps to visualize genomic interactions on the final primary assemblies of each strain. To those ends, we aligned paired-end Hi-C reads of each strain to each scaffolded assembly using BWA-MEM [91]. We created a normalized contact matrix with the aligned reads using *HiCExplorer* version 3.7.2 [92]. We used *HiCExperiment* and *HiContacts* (Bioconductor) packages to visualize Hi-C contact maps for genome assemblies [93].

### Genome annotation

Prior to annotation, assembled scaffolds were screened for taxonomic contamination using FCS-GX (Foreign Contamination Screen - Genome eXclusion, https://github.com/ncbi/fcs) [94]. Structural and functional genome annotation was then conducted using EGAPx (Eukaryotic Genome Annotation Pipeline – External, https://github.com/ncbi/egapx). See details in supplemental methods in additional file 1.

### Chromosomal synteny analysis

To assess chromosomal-level conservation and syntenic relationships, we employed GENESPACE following the published protocol [95]. Annotated gene sets from the genome assemblies of each *M. domestica* strain, *D. melanogaster* (GCF_000001215.4), and *L. cuprina* (GCF_022045245.1) were input into GENESPACE. Orthologous genes were identified across genomes using sequence similarity clustering, and genomic positions were used to define syntenic blocks (5 genes in a row). GENESPACE then generated a synteny map and visualization output, enabling comparison of synteny, chromosomal rearrangements, and conserved gene order. *D*. *melanogaster* was used as a reference to establish chromosomal assignments.

### Assignment of homologous genes between *D. melagnogaster*, *L. cuprina*, and house fly strains

We used peptide sequences of the protein-coding genes from *D. melanogaster*, *L. cuprina*, or house fly strains as queries in *BLASTP* searches within the option ‘-evalue 1e-5’ (*BLAST+* version 2.13 [96]) against proteins of each other species, retaining only the target hit with the top bit score for each query. We also used these best hits to perform reciprocal best *BLASTP* searches.

### Dot plot alignment visualization

Assemblies were aligned with the *nucmer* aligner from the *MUMmer4* package [97] using an alignment seed length of 2000 bp and the coordinates output format. Assembly fasta files were indexed with *Samtools* version 1.17 [98]. The coordinates and fasta index files were then plotted using custom R and ggplot2 scripts [99] available at https://github.com/jaehakson/houseflygenomes.

### Sequencing coverage of chromosomes

We used *Minimap2* version 2.26 to generate the alignment file of PacBio HiFi raw sequencing reads against the primary genome assemblies [100], and we then filtered the alignment with a minimum mapping quality ‘20’ using *Samtools* version 1.18 [98]. We used *Bedtools* to measure the sequencing coverage in a 10,000bp window and calculated the median of the sequencing coverage in 10kbp windows across chromosomes I-V, excluding chromosomes X or Y. We next divided the coverage of each window by the genome-wide (chromosomes I-V) median of the sequencing coverage to normalize the sequencing coverage of each window. We used only windows from chromosomes I-V to determine the denominator used for normalization for two reasons. First, the house fly X or Y chromosomes are highly heterochromatic [44,101], and so our analysis of filtering out the reads with low mapping quality could underestimate the sequencing coverage on the X or Y. Second, the house fly MY strain has one copy of X and Y chromosomes, indicating the half in its sequencing coverage relative to other chromosomes.

Through this normalization method, we recalibrated the coverage of autosomes to one, which denotes two copies of homologous chromosomes in the genome for the mapping features.

Tandem repeat visualization within the pericentric regions of the proto-Y chromosomes.

We used *ModDotPlot* version 0.9.9 [102] to visualize the complex repeats and repeat arrays within the pericentric regions, including the centromeres, of the *Mdmd*-bearing proto-Y chromosomes.

### *k*-mer approach for Y chromosome Genome Scan (YGS)

We took a *k-mer* comparison approach, called YGS, [103] to test enrichment of female sequences on the Y chromosome, and the other chromosomes at the contig level. In the YGS run, we used a *k*-mer size of 15bp, used the same strain male sequencing reads for a validating bit-array, and employed the same options used in the *Drosophila* example from [103].

### Divergence in protein-coding and non-coding sequences

We lifted over the NCBI RefSeq annotation of the house fly genome (GCF_030504385.1) to each haplotype assembly separately using *Liftoff* version 1.6.3 [104], and extracted protein-coding sequences with each lifted-over annotation using *GffRead* version 0.12.7 [105]. To align the protein-coding sequences between two haplotypes, we used a program ‘alignSequences’ from *MACSE* version 2.07 [106]. We then used the other program ‘reportMaskAA2NT’ from *MACSE* to perform a post-filtering of the protein-coding sequence alignment for downstream analyses. The final alignments between two alleles were used to calculate the rate of nonsynonymous (*K_a_*) and synonymous (*K_s_*) substitutions using *KaKs_Calculator 3.0* [107].

We first aligned PacBio HiFi reads of each strain’s males to the corresponding strain genomes with *Minimap2* version 2.26 [100], and then converted to BAM files with *Samtools* version 1.18 [98]. To call variants, we ran *Deepvariant* version 1.9.0 with the parameter ‘-- model_type=PACBIO’ with the BAM files of each strain’s males [108]. We selected variants with ‘PASS’ in the filter column and the heterozygous genotype from the output of *Deepvariant*. We used *VCFtools* version 0.1.16 to calculate the heterozygous variant density in the 1Mb window.

### Comparing heterozygosity across chromosomes and strains

We used a modified version of Tajima’s *D* statistic [109] to compare heterozygosity on each chromosome across the strains. We first identified the fraction of synonymous sites that differ between haplotypes, within protein-coding sequences for each strain. We then calculated a modified Tajima’s *D* statistic (D_kab_) between the MY strain and each of the other strains within the moving average of 50 genes on each chromosome, using the filter function in R [99]. See details in supplemental methods in additional file 1.

### Repeat annotation

Repetitive elements in the genome assemblies were identified, curated, and annotated across the unmasked genome assembly using the fully automated Earl Grey pipeline (version 7.3.10) [110]. See details in supplemental methods in additional file 1.

### Phylogenetic tree of *Mdmd* and Ncm

To generate a phylogenetic tree of Mdmd and Ncm (Nucampholin) of other flies, we employed Ncm (a homolog of CWC22 spliceosome-associated protein) from *Musca vetustissima* (XP_061386924.1), *S. calcitrans* (XP_013100431.2), H. irritans (XP_075152361.1), *C. vicina* (XP_065358704.1), and *L. cuprina* (XP_023307808.2) and we used Mdmd and Ncm of each house fly strain obtained from each house fly assembly genome. We then aligned amino acid sequences with MUSCLE version 5.1.0 [111] and trimmed the alignment with trimAl version 1.5 with a parameter ‘-gt 0.5’ [112]. This final alignment was used as input when running IQ-TREE version 2.3.6 with parameters ‘-m MFP -B 1000’ to obtain the bootstrap support in the tree construction [113,114].

### *M* locus comparisons

Using *Minimap2* version 2.26 [100], we performed pairwise comparisons among II^M^, III^M^, and Y^M^ chromosomes to examine the *M* locus in each strain. We kept matches with mapping quality of more than ‘60’ in the PAF format, and then we implemented the number of residue matches and the alignment block length to calculate the sequence identity among the *M* loci of the three (M2, M3, and MY) strains. We also identified the identities in coding sequences near the *M* locus between the M3, and MY strains, using BLAST+ version 2.13 in the BLASTN search [96]. We used the *‘circlize’* package in R to visualize *M* locus among the three strains [115].

## Supporting information

Additional file4: Phylogenetic tree for Mdmd and Ncm of flies in a Newick format

## Acknowledgements

Computational analyses were performed on the Carya cluster in the Research Computing Data Core (RCDC) at the University of Houston, on the Amarel cluster in the Office of Advanced Research Computing (OARC) at Rutgers, The State University of New Jersey, and on computing resources provided by the SCINet project and/or the AI Center of Excellence of the USDA Agricultural Research Service. This study was supported by the USDA Agricultural Research Service with the project “Exploiting Veterinary Pest Genomics to Improve Animal Health” (3094-10400-004-000-D).

## Availability of data and materials

Data was deposited into NCBI’s Sequence Read Archive (SRA) and genome assemblies of each strain are available under BioProject PRJNA1504915. Sample accessions for raw sequencing data are retained as: MY-aabys, SAMN62070911; M1-aabys, SAMN62070912; M2, SAMN62070913; M3, SAMN62070914; M5, SAMN62070915. Analysis code can be found in the following GitHub repository: https://github.com/jaehakson/houseflygenomes

## Supplementary Information

Additional file1: Supplemental Figures; Figure S1 to S9

Additional file2: Supplemental Tables

Table S1. Number of house fly genes that are a best hit and not a best hit of L. cuprina X genes (Chi-squared test).

Table S2. *k*-mer matches of MY male assembly (denoting chromosome and contigs in the separate columns).

Table S3. *K_s_* values between two haplotypes for each strain Table S4. Heterozygous variant density

Table S5. Interproscan output with six peptide sequences that are shared in *M*-adjacent regions between the M3 and MY strains.

Table S6. House fly strains and summary of sequencing process methodology Additional file3: Supplemental Text and Supplemental Methods

Additional file4: Phylogenetic tree for Mdmd and Ncm of flies in a Newick format

