## Additional file1: Supplemental Figures for "X-Y divergence of house fly (*Musca domestica*) proto-sex chromosomes follow distinct evolutionary trajectories despite residing in the same genome"

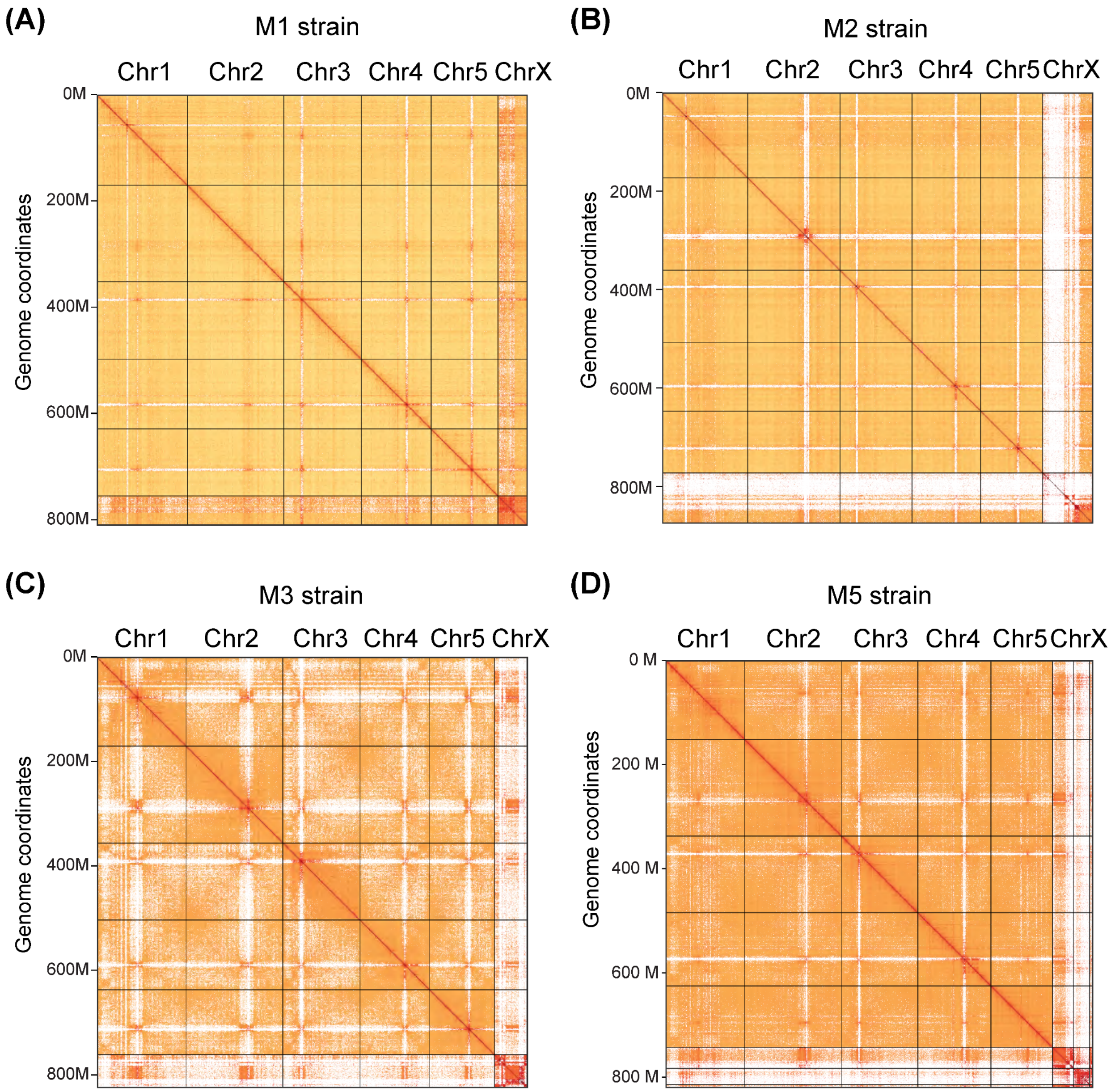


Figure S1. HiC contact map from six chromosomes (I-V, and X) of the M1 (A), M2 (B), M3 (C), and M5 (D) strains

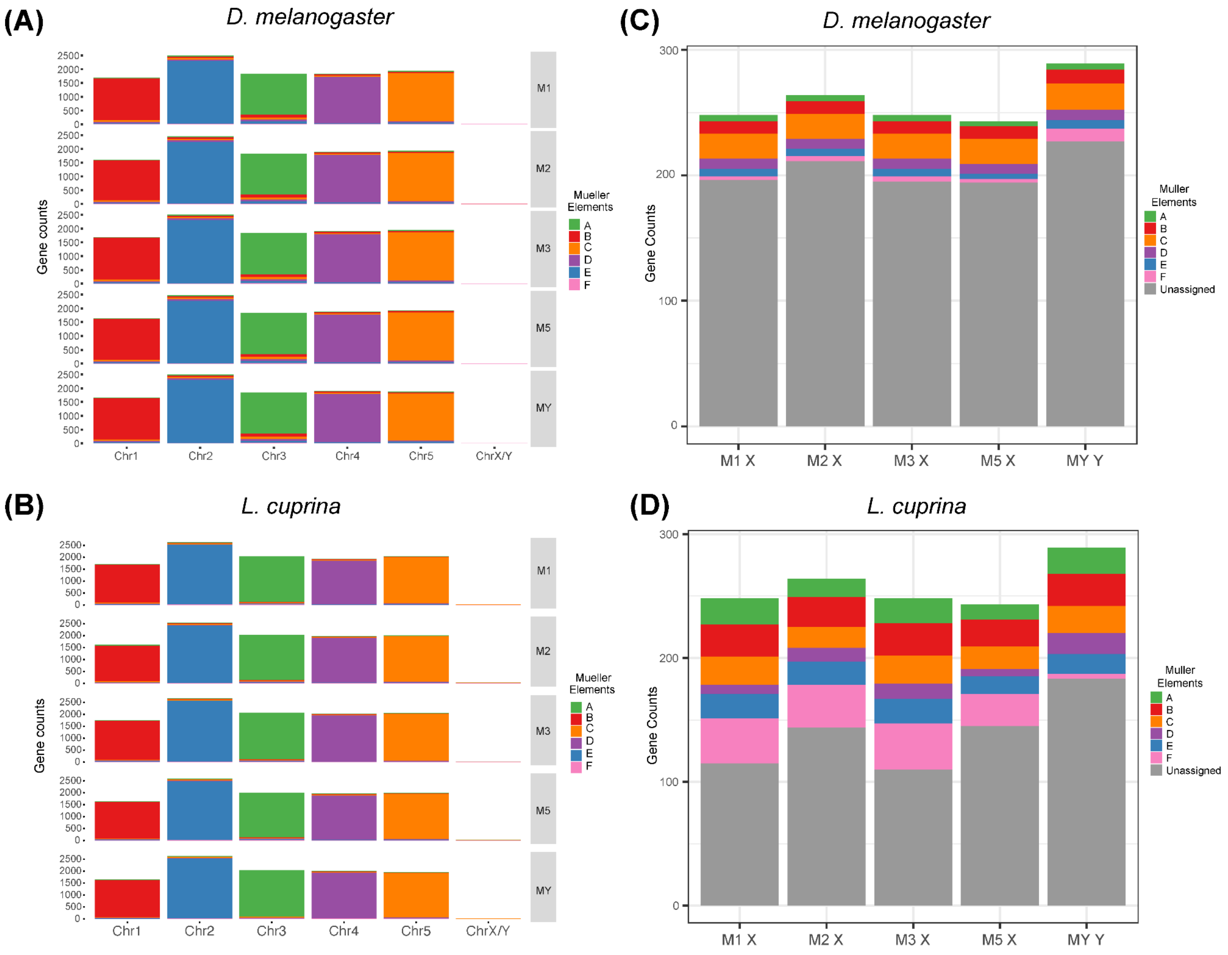


Figure S2. Protein-coding gene mapping by reciprocal (A, B) and one-way (C, D) best-matched blast hits between house fly strains and *D. melanogaster* and *L. cuprina*. (A) House flies vs *D. melanogaster*, (B) House flies vs *L. cuprina*, (C) House fly X genes to *D. melanogaster* genome, (D) House fly X genes to *L. cuprina* genome

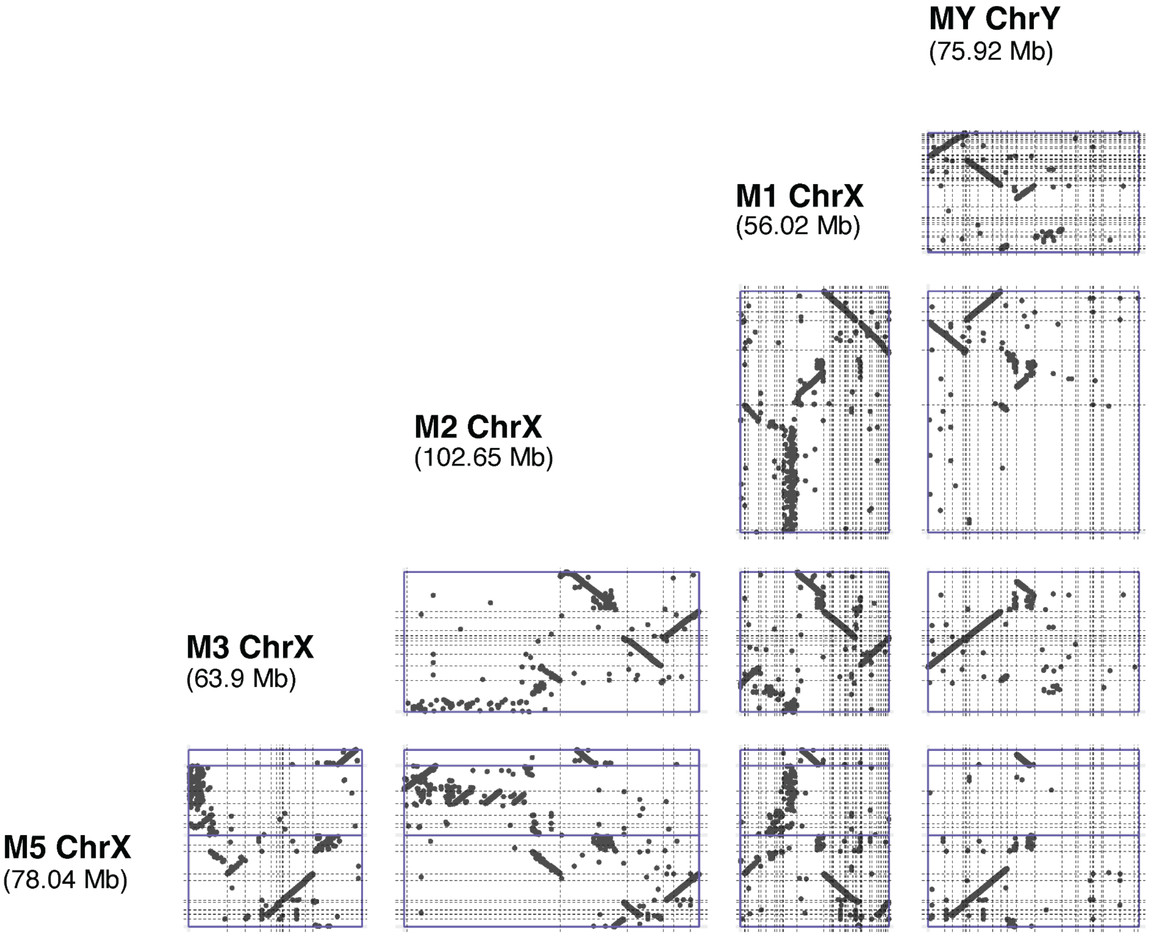


Figure S3. Dotplots of the X/Y chromosomes across all five (MY, M1, M2, M3, and M5) strains in all pairwise comparisons

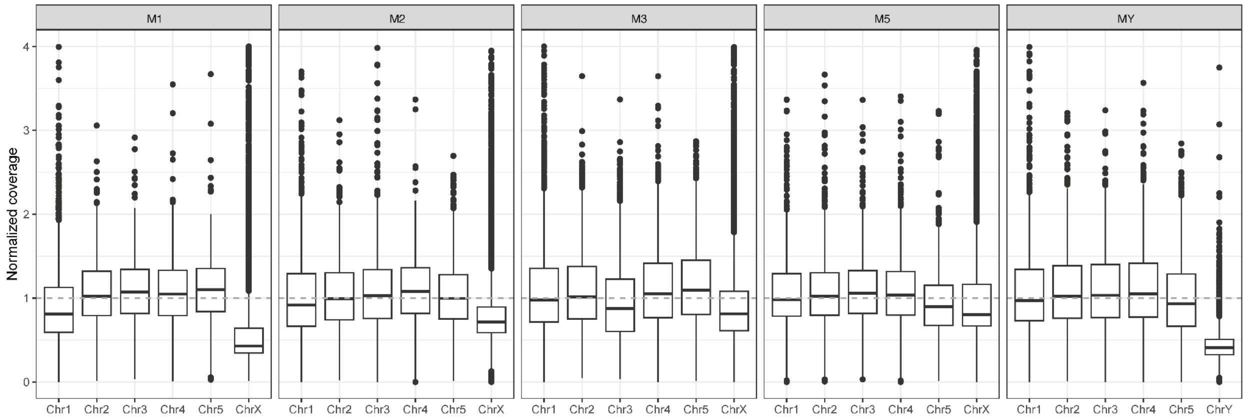


Figure S4. Box plots that show normalized coverage of male reads to the chromosomes in the genome assembly of the strain from which the reads were sampled

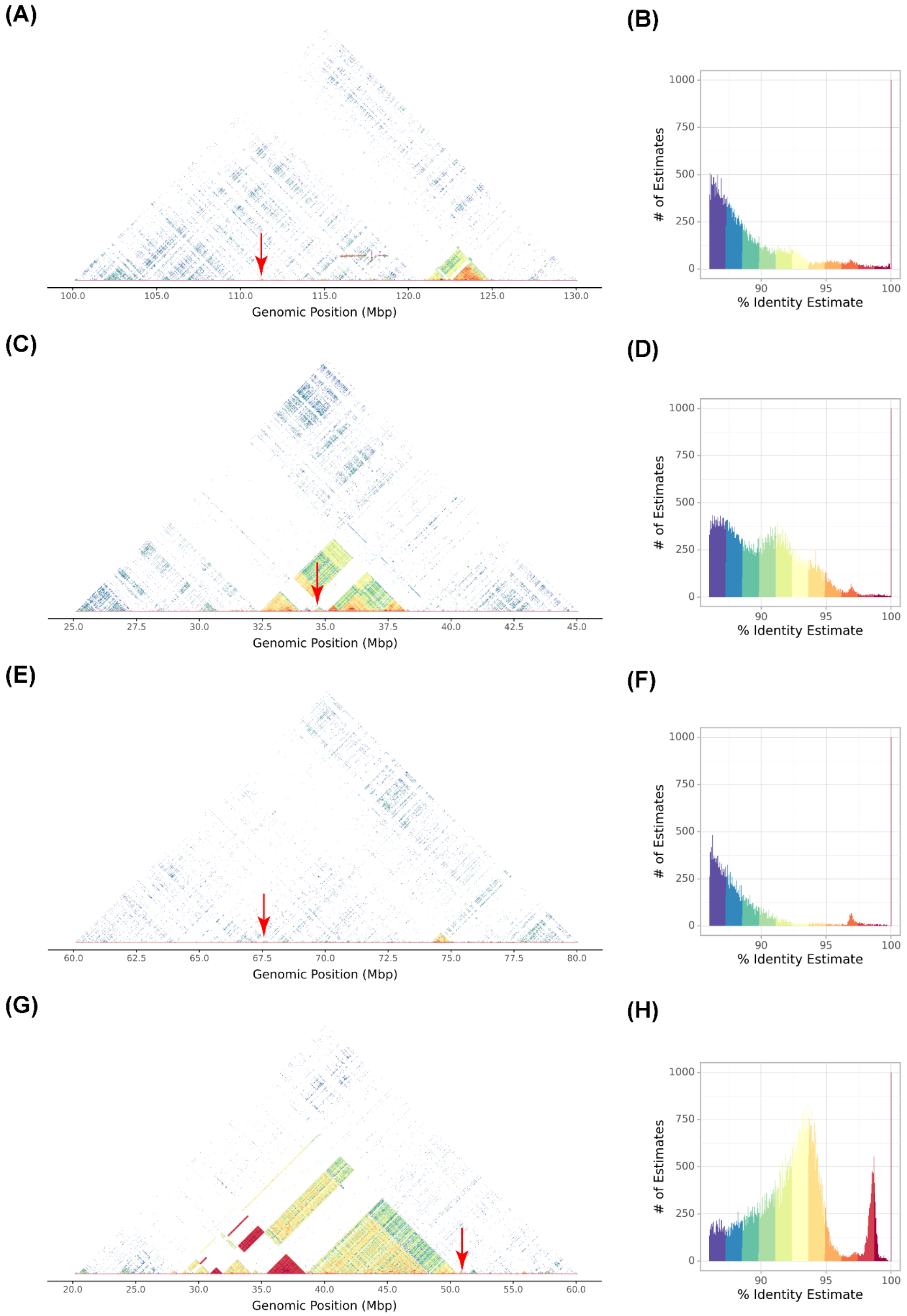


Figure S5. Visualization of tandem repeats in pericentric regions of the proto-Y chromosomes. Diagonal self-identity heatmap of M2 (A), M3 (C), M5 (E), and MY (G) strains and the estimated percent identity and number of alignments of M2 (B), M3 (D), M5 (F), and MY (H) strains. High percent identities of tandem repeats (>90%) are inferred as centromeric regions of the chromosomes.

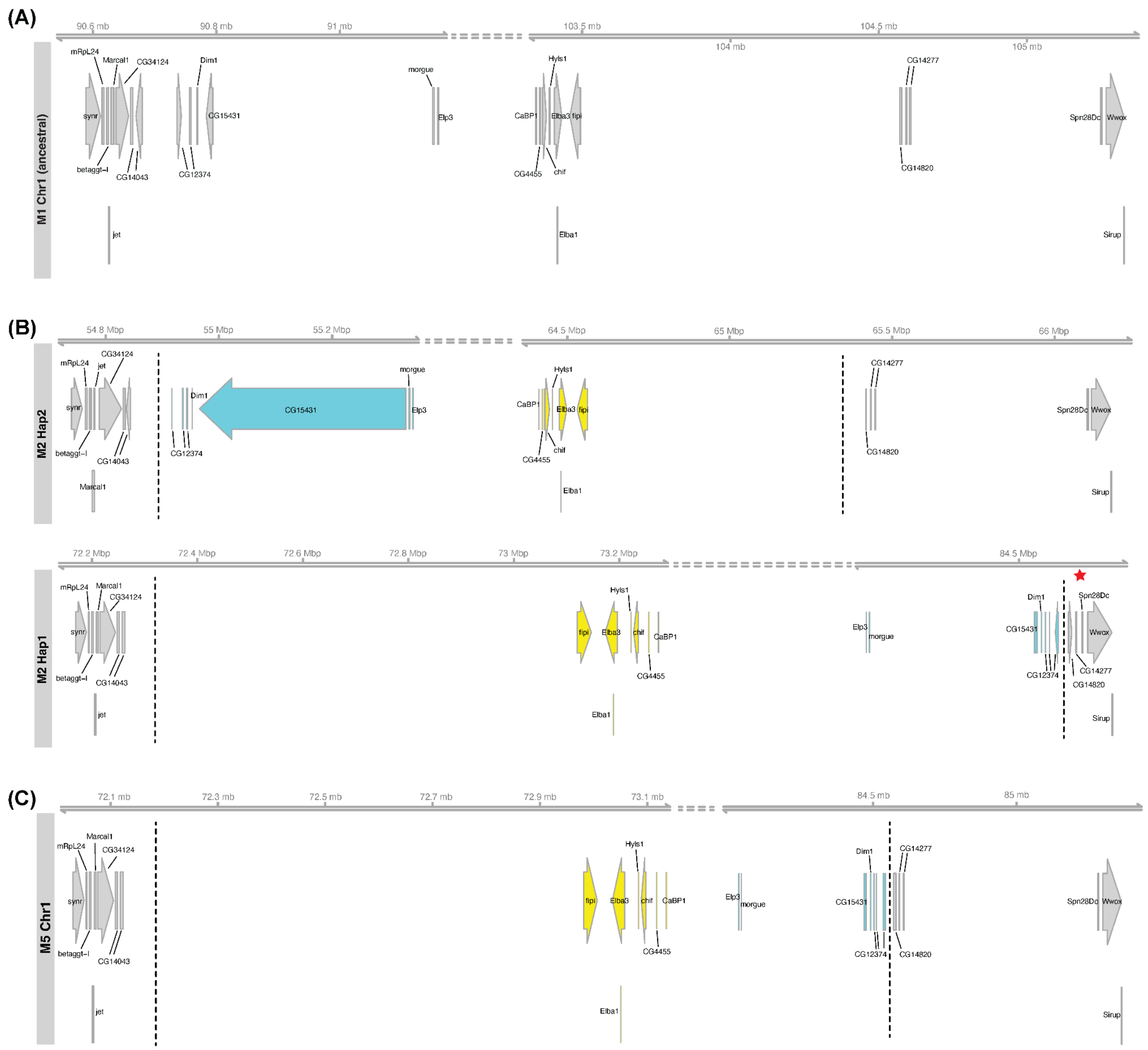


Figure S6. Synteny around chromosome I inversion breakpoints from the M1(A), M2 (B) and M5 (C) strains; *D. melanogaster* gene symbols, homologous to the house fly genes, were used. As the M2 strain has the heterozygous inversion, synteny from haplotype 1 and 2 is displayed in (B). Red stars represent the locations of assembly gaps on scaffolds.

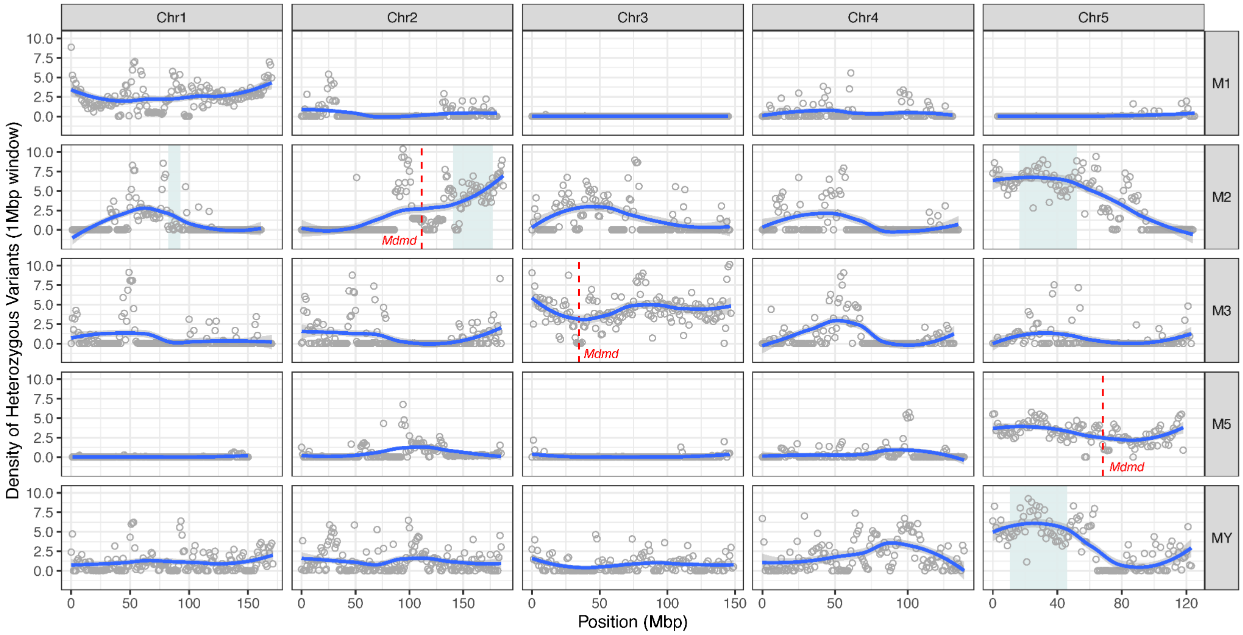


Figure S7. Density of heterozygous variants across chromosomes of the house fly strains. Blue smooth lines across each chromosome are presented for the heterozygous variant density points (1Mbp window). Cyan shaded regions indicate polymorphic (heterozygous) inversions in the M2 and MY strains.

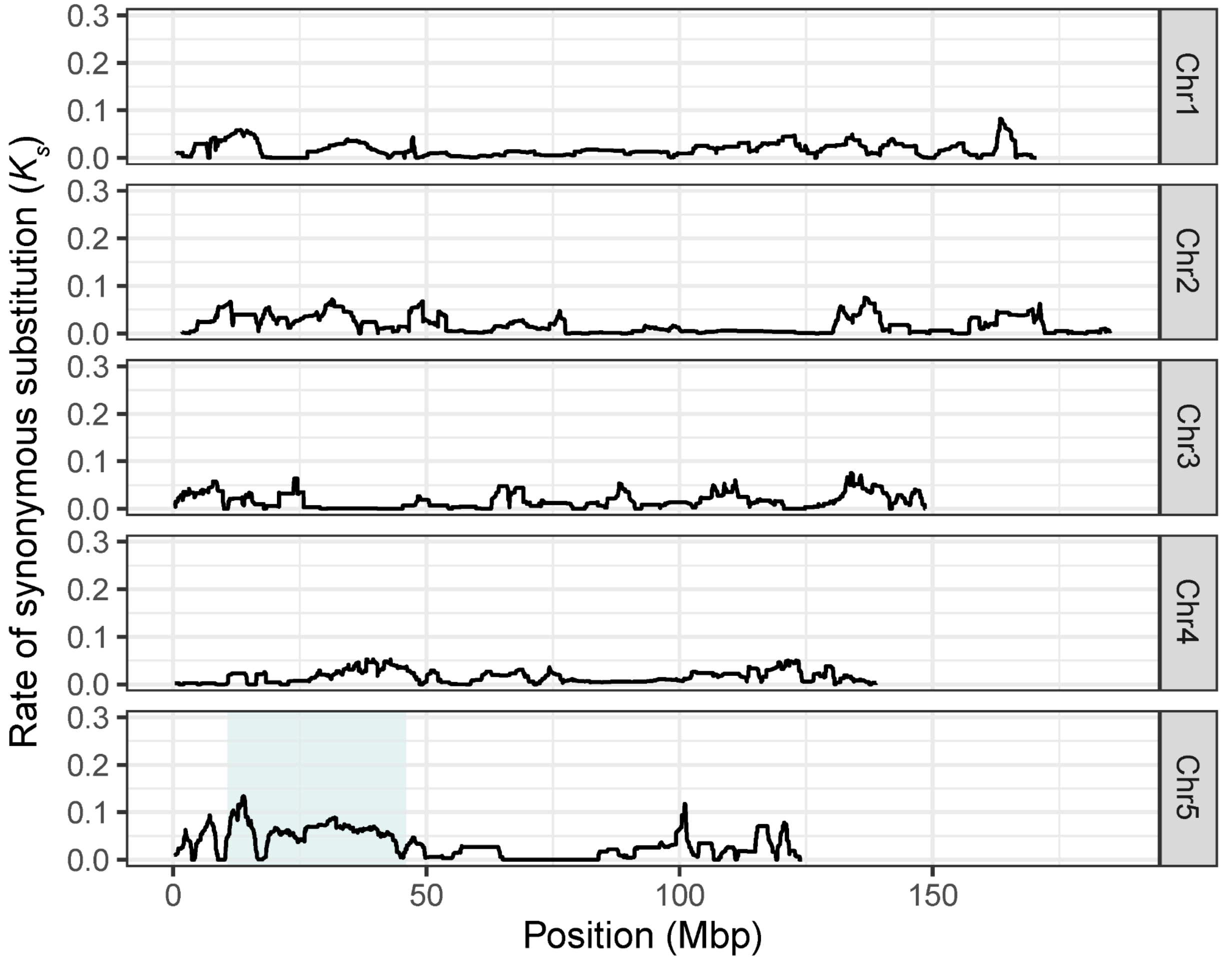


Figure S8. Synonymous divergence ($K_s$) between the homologous chromosomes of the MY strain. Cyan shaded region in chromosome 5 indicates a heterozygous inversion in the MY strain.

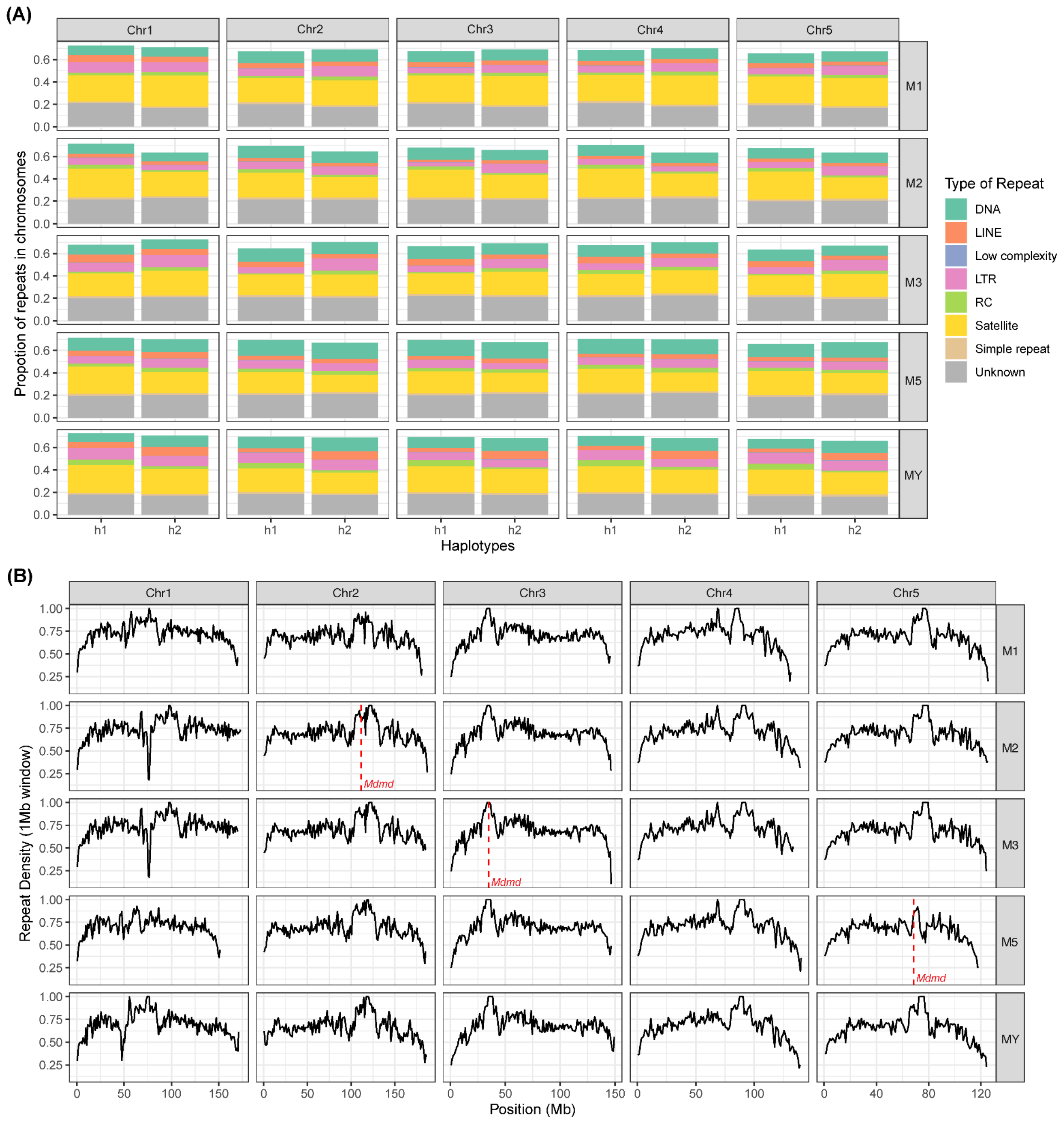


Figure S9. (A) Repeat compositions in each of two homologous chromosomes (two haplotype assemblies) of the house fly strains. (B) Repeat density across chromosomes of the house fly strains in primary assemblies.
