## Additional file3: Supplemental Text and Supplemental Methods for "X-Y divergence of house fly (*Musca domestica*) proto-sex chromosomes follow distinct evolutionary trajectories despite residing in the same genome"

Best blast hits between house fly strains and *L. cuprina*

We found that 25-30% of annotated genes on the house fly X or $Y^M$ chromosome in each strain had their best hit on the *L. cuprina* X chromosome (Figure 3C-F). However, most of those house fly genes hit the same *L. cuprina* gene (LOC124418751), which encodes a protein with a high sequence identity to a predicted protein from a Wolbachia endosymbiont (DUF1759). This protein is part of a family of gag-proteins (PF03564), suggesting that it is of retroviral origin. There were only 4-6 other *L. cuprina* X-linked genes that were best hits, each with 2-10 matches on the house fly X chromosome.

### Supplemental methods

PacBio HiFi Sequencing

DNA was extracted from a single unfed, adult, male individual from each strain. Each sample was homogenized in a 1.5 mL tube in a liquid nitrogen bath with a liquid nitrogen-cooled pestle. The Qiagen MagAttract HMW DNA Kit (Qiagen, Hilden, Germany) was used to obtain sufficiently high molecular weight (HMW) DNA (Table S6). Immediately following homogenization, 200 µl 1X PBS, 20 µl Proteinase K, 8 µl RNase A, and 150 µl Buffer AL were added to the sample. The tube was slowly inverted to mix. Samples were incubated for two hours at 25°C and 900 rpm. Lysate was transferred to a 2.0 mL tube using a wide-bore pipette tip following brief centrifugation. Qiagen's fresh or frozen tissue protocol was followed starting with the addition of Buffer MB. Modifications to the protocol were as follows. All washes and rinses were incubated for 1 minute. DNA was eluted with 105 Buffer AE and incubated at room temperature for 5 minutes at 1400 rpm. Following isolation, genomic DNA was quantified using the Broad Range dsDNA Qubit assay (Thermo Fisher Scientific, Waltham, MA, USA). Size distribution was assessed with the Genomic DNA 165 kb kit run on the Femto Pulse System (Agilent Technologies, Santa Clara, CA, USA). A PacBio SMRTbell library was prepared using the SMRTbell Express Template Prep Kit 2.0 and associated guide for preparing whole genome libraries (Pacific Biosciences, Menlo Park, CA, USA). The prepared library was bound and sequenced at the USDA-ARS Veterinary Pest Genetics Research Unit in Kerrville, Texas, on Pacific Biosciences 8M SMRT Cells with a Revio system (Pacific Biosciences, Menlo Park, CA, USA) beginning with a 2-h pre-extension followed by a 30-h movie collection time. After sequencing, circular consensus sequences from the PacBio Revio subreads were obtained using the SMRTLink v13.0 software. For the MY strain, we performed HMW DNA extraction with three males using the Monarch HMW DNA Extraction Kit (T3060) from New England Biolabs (NEB) following its tissue protocol (Table S6). Three separate HMW DNA extractions were performed and sent to Novogene Corporation Inc. They prepared Pacbio sequel II DNA HiFi library for each DNA extraction for being bound and sequenced on Pacbio sequel II (HiFi/CCS mode/cell).

Hi-C Library preparation

Hi-C library was generated using the Phase Genomics Proximo Animal kit version 4.0. Approximately 500 mg tissue was finely chopped and then crosslinked for 15 minutes at room temperature with end-over-end mixing in 1 mL of Proximo crosslinking solution. Crosslinking reaction was terminated with a quenching solution for 20 minutes at room temperature again with end-over-end mixing. Quenched tissue was rinsed once with 1X Chromatin Rinse Buffer (CRB). Tissue was transferred to a liquid nitrogen-cooled mortar and ground to a fine powder. Powder was resuspended in 700 µL Proximo Lysis Buffer 1 and incubated for 20 minutes with end-over-end mixing. A low-speed spin was used to clear the large debris and the chromatin containing supernatant transferred to a new tube. Following a second higher speed spin, the supernatant was removed and the pellet containing the nuclear fraction of the lysate was washed with 1X CRB. After removing 1X CRB wash, the pellet was resuspended in 100 µL Proximo Lysis Buffer 2 and incubated at 65°C for 15 minutes. Chromatin was bound to Recovery Beads for 10 minutes at room temperature, placed on a magnetic stand, and washed with 200 µL of 1X CRB.

Chromatin bound on beads was resuspended in 150 µL of Proximo fragmentation buffer and 2.5 µL of Proximo fragmentation enzyme added and incubated for 1 hour at 37°C. Reaction was cooled to 12°C and incubated with 2.5 µL of finishing enzyme for 30 minutes. Following the addition of 6 µL of Stop Solution, the beads were washed with 1X CRB and resuspended in 100 µL of Proximo Ligation Buffer supplemented with 5 µL of Proximity ligation enzyme. Proximity ligation reaction was incubated at room temperature for 4 hours with end-over-end mixing. To this volume, 5 µL of Reverse Crosslinks enzyme was added and the reaction incubated at 65°C for 1 hour.

After reversing crosslinks, the free DNA was purified with Recovery Beads and Hi-C junctions were bound to streptavidin beads and washed to remove unbound DNA. Washed beads were used to prepare paired-end deep sequencing libraries using the Proximo Library preparation reagents.

Library quality control and sequencing

Library fragment size and concentration were determined using TapeStation capillary electrophoresis and quantitative PCR. Prior to production sequencing, library performance was determined by low-pass sequencing, generating ~200,000 read pairs of sequencing data on an Illumina iSeq System. Resulting reads are mapped to a reference assembly and analyzed with Phase Genomics' open-source QC tool, hic_qc (GitHub - phasegenomics/hic_qc: A (very) simple script to QC Hi-C data). Libraries receiving a 'Sufficient' quality judgement from hic_qc are advanced to production sequencing. Samples were sequenced on a NovaSeq X, 300 cycles, PE150.

Genome annotation and chromosomal synteny analysis

FCS-GX compares assembly fragments against a curated reference database and flags scaffolds with strong similarity to non-target taxa; scaffolds identified as likely contaminants were removed or manually inspected following standard filtering thresholds. Only retained scaffolds were used for downstream annotation.

EGAPx integrates repeat masking, transcript and protein evidence, RNA-seq data, and ab initio prediction. Strain specific transcriptomic reads were provided to EGAPx where available for the transcript assembly and alignment part of the pipeline. EGAPx further utilizes protein homology evidence from related taxa to provide additional exon and coding-region support. The final EGAPx annotation is a combination of transcript and protein-based evidence with ab initio predictions to produce consensus gene models optimized for completeness and structural accuracy. The functional annotation assigns protein domains, orthology information, and Gene Ontology terms, resulting in a comprehensive gene set including protein-coding genes, noncoding RNAs, pseudogenes, and transposable elements. The genome assemblies including the primary and the haplotypes were annotated separately.

Comparing heterozygosity across chromosomes and strains

From each diploid phased genome assembly, $\pi_s$ was calculated as the fraction of synonymous sites that differ between haplotypes, within protein-coding sequences. We interpret $\pi_s$ as an estimator of $\Theta = 4N\mu$, referred to as $\hat{\Theta}_\pi$ .

Tajima (Tajima 1989) showed that we can compare $\hat{\Theta}_\pi$ with $\Theta$ estimated using the number of segregating sites, or $\hat{\Theta}_W$ (Watterson 1975). Tajima's D statistic is the measure of the deviation between the estimate of $\Theta$ from the number of pairwise difference ($\hat{k}$) and the estimate from the number of segregating sites ($S$):

$$D = \frac{d}{\sqrt{\hat{V}(d)}} = \frac{\hat{k} - \frac{S}{a_1}}{\sqrt{\hat{V}(d)}}$$

We modified this calculation as follows. Instead of comparing $\hat{\Theta}_\pi$ and $\hat{\Theta}_W$, we compared $\hat{\Theta}_\pi$ (i.e., $\hat{k}$) estimated from two different samples. In our case, the two samples are the estimates of $\hat{\Theta}_\pi$ from two different haploid assemblies from two separate strains. To do so, we defined the pairwise differences (per site within a window length $l$) between haplotypes (i.e., $\pi_s$) in strain $a$ as $\hat{k}_a$, and, likewise, between haplotypes in strain $b$ as $\hat{k}_b$. This allows us to write the differences $\hat{k}$ between strains as:

$$d = \hat{k}_a - \hat{k}_b$$

The denominator of Tajima's $D$ statistic includes the sample variance of $d$, or $\hat{V}(d)$. In our case, $\hat{V}(d) = V(\hat{k}_a) + V(\hat{k}_b)$. Tajima showed that:

$$V(\hat{k}_a) = b_1\Theta + b_2\Theta^2,$$

where

$$b_1 = \frac{n+1}{3(n+1)},$$

and

$$b_2 = \frac{2(n^2+n+3)}{9n(n-1)}.$$

In the case of comparing two haplotypes, $n$ = 2, which gives us $b_1 = 1$ and $b_2 = 1$. Therefore,

$$V(\hat{k}_a) = \hat{k}_a + \hat{k}_a^2$$

and

$$V(\hat{k}_b) = \hat{k}_b + \hat{k}_b^2$$

because $\hat{k}$ is an unbiased estimator of $\Theta$. We can therefore write our modified version of Tajima's $D$ as:

$$D_{kab} = \frac{\hat{k}_a - \hat{k}_b}{\sqrt{(\hat{k}_a + \hat{k}_a^2) + (\hat{k}_b + \hat{k}_b^2)}}.$$

Repeat annotation

Known repeat elements were initially identified by querying the assemblies against the Dfam 4.0 database via FamDB using RepeatMasker (v4.2.4) (Smit et al. 2013). Unannotated *de novo* repeat families were subsequently identified using RepeatModeler2 (v2.0.9) (Flynn et al. 2020), incorporating LTRfinder (Xu and Wang 2007)to detect full-length LTR retrotransposons. Raw *de novo* models were refined and extended through Earl Grey's automated "BLAST, Extract, Align, Trim" (BEAT) iterative framework to better define boundary sequences, followed by clustering to eliminate consensus redundancy. The repeat annotations from the output were further used for downstream analyses and statistical comparisons.

Supporting data 1. Normalized coverage of each strain
